# Genome evolution at the extreme of angiosperm miniaturization

**DOI:** 10.64898/2026.08.20.746096

**Authors:** Aidi Zhang, Zhongyu Tang, Na Wei

**Author notes:** These authors contributed equally to this work.

## Abstract

Eukaryotic genomes vary by several orders of magnitude, yet this vast variation bears little relation to the complexity of the organisms they encode. Thus, how genome evolution accompanies changes in organismal complexity remains unresolved. A central obstacle is that major differences in body plan usually occur among deeply divergent lineages, entangling body plan evolution with the genomic divergence accumulated over long independent histories. Duckweeds offer a rare system in which successive body plan reductions can be traced within a single plant family. Across this trajectory, body size declined by nearly an order of magnitude and roots were progressively lost, culminating in the extreme of angiosperm miniaturization. Yet genome size increased nearly sixfold. Here, using a new chromosome-scale genome of *Wolffia globosa* and comparative genomics across nested evolutionary scales, we show that genome size, gene number, and functional repertoire followed distinct trajectories during miniaturization. Genome expansion was driven largely by transposable element accumulation, whereas the number of protein-coding genes remained stable. Aquatic adaptation itself promotes functional simplification, but establishes only a baseline. Duckweeds pushed this streamlining much further through additional contraction of developmental, structural, and biotic defense functions, alongside selective expansion of functions associated with growth and abiotic adaptation. This remodeling accumulated across successive evolutionary transitions through continued contraction of the same gene families and, more commonly, contraction of different families affecting the same biological processes. Organismal complexity may therefore reflect not simply the size of a genome or its functional repertoire, but how that repertoire is selectively reconfigured through evolution.

## Introduction

Throughout evolution, organisms have repeatedly evolved diverse and complex body plans. Examples include the emergence of multicellular life, the colonization of land by plants and animals, and the diversification of specialized tissues and organs across major lineages (Donoghue et al., 2021; Niklas & Newman, 2013; Wei et al., 2026). These evolutionary transitions are accompanied by changes in genome architecture, suggesting a correspondence between genomic evolution and changes in organismal complexity. Because greater organismal complexity was thought to require more genetic instructions to encode increasingly elaborate biological organization, complex organisms were historically expected to possess larger genomes (Gregory, 2002; Gregory, 2005). At broad evolutionary scales, genome expansion appears to accompany major increases in biological organization: prokaryotes generally possess compact genomes, whereas eukaryotes typically have larger genomes, with multicellular lineages often exceeding unicellular lineages. Yet the relationship between genome evolution and organismal complexity remains surprisingly elusive (Alvarez-Ponce & Krishnamurthy, 2025). The C-value, defined as the amount of nuclear DNA in a haploid genome, varies by several orders of magnitude across eukaryotes with little consistent correspondence to organismal complexity (Gregory, 2005). This apparent disconnect challenged the assumption that genome expansion parallels organismal complexity and lies at the heart of the “C-value paradox” (Elliott & Gregory, 2015; Gregory, 2005). The paradox raises a fundamental question: how does genome evolution accompany changes in organismal complexity?

Genome size captures only one dimension of genome evolution. The accumulation of non-coding and repetitive DNA explains why genome size can vary independently of protein-coding gene content (Elliott & Gregory, 2015). Gene number itself also shows surprisingly little correspondence with organismal complexity, giving rise to the “G-value paradox” (Hahn & Wray, 2002). Thus, neither genome size nor gene number alone provides a sufficient explanation for how genomic changes accompany organismal complexity. Different evolutionary processes can further reshape these genomic dimensions in distinct ways. Transposable element (TE) proliferation can dramatically expand genomes without proportional increases in protein-coding genes and represents a major driver of genome size variation in plants (Bao et al., 2026; Bennetzen et al., 2005). Gene duplication and whole-genome duplication (WGD) generate paralogs that can be retained, lost, or functionally diversified, providing important substrates for evolutionary innovation (Soltis & Soltis, 2016; Van de Peer et al., 2017). Such differential gains and losses can reshape gene family repertoires and alter the functional potential encoded by genomes. It has been proposed that gene family diversity and composition may provide additional insights into organismal complexity beyond genome size and gene number (Vogel & Chothia, 2006). Although this hypothesis has been explored using cell type number as a proxy for complexity, it remains largely untested across evolutionary transitions in organismal form (Alvarez-Ponce & Krishnamurthy, 2025). Together, genome size, gene number, and functional repertoire capture distinct but interconnected dimensions of genome evolution.

Determining how genomic dimensions evolve with organismal complexity is challenging because major differences in body plan often occur among deeply divergent lineages. Extensive genomic divergence can obscure the genomic changes associated with body plan evolution. Duckweeds (Araceae, subfamily Lemnoideae) provide a remarkable exception. The transition from the relatively large, multi-rooted *Spirodela* through the smaller, single-rooted *Lemna* to the highly reduced *Wolffia* (Fig. S1), provides a natural evolutionary series of progressive body plan reduction. Along this trajectory, body size and vegetative architecture became progressively reduced, accompanied by the reduction or loss of roots, conventional leaves and stems, and vascular differentiation (Acosta et al., 2021; Ware et al., 2023). These successive reductions culminated in the rootless *Wolffia*, the smallest flowering plants and one of the most extreme simplifications of the angiosperm body plan (Fig. S1). Remarkably, duckweeds present a striking evolutionary contrast: progressive simplification of the body plan occurred alongside expansion of the genome (Hoang et al., 2022). When and how different genomic dimensions evolved along this trajectory, however, remain unknown. Yet this trajectory toward miniaturization evolved in the context of a return from terrestrial to aquatic life.

The transition from terrestrial to aquatic life has occurred repeatedly across vascular plants, particularly within angiosperms, enabling recurrent genomic features of aquatic adaptation to be identified across diverse lineages (Bowles et al., 2022; Chen et al., 2026; Guo et al., 2025). Comparative genomic studies across phylogenetically diverse aquatic plants have identified recurrent gene losses and gene family contractions, together with lineage-specific expansions, affecting functions such as cell wall biosynthesis, lignification, hypoxia responses, stomatal development, immunity, and root-associated pathways (Guo et al., 2025; Meseguer et al., 2022). Together, these changes point to convergent remodeling of functions associated with structural support, gas exchange, environmental responses, and organ development during the transition to aquatic life. Yet extreme miniaturization is not a general outcome of this transition. Most aquatic plants retain substantially greater body size and structural complexity than duckweeds. Among aquatic growth forms, free-floating and floating-leaved plants provide a particularly relevant context for understanding duckweed evolution. They share the air–water interface occupied by duckweeds while spanning a broad spectrum of body size and structural complexity, from rooted species with complex floating leaves to free-floating plants with reduced vegetative bodies (Chen et al., 2026). Duckweeds represent the extreme endpoint of this spectrum. Distinguishing genomic changes accompanying this extreme miniaturization from those broadly associated with aquatic adaptation is therefore essential.

To understand how genome evolution accompanied progressive body plan simplification in duckweeds, we combined chromosome-scale genomics with comparative evolutionary analyses across nested evolutionary scales. We generated a chromosome-scale genome of *Wolffia globosa*, extending genomic representation of the *Wolffia* lineage beyond a single species (Michael et al., 2021). This enabled lineage-level reconstruction of genomic evolution toward extreme miniaturization. We then asked whether genome size, gene number, and functional repertoire followed similar or distinct evolutionary trajectories as the duckweed body plan became progressively simplified. Comparisons across phylogenetically diverse aquatic plants placed these changes in the context of the recurrent transition to aquatic life, allowing us to distinguish patterns broadly associated with aquatic adaptation from those associated with extreme miniaturization. Finally, we reconstructed gene family evolution across duckweeds to determine how functional repertoires changed along the trajectory from *Spirodela* through *Lemna* to *Wolffia* lineages (Acosta et al., 2021; Ernst et al., 2025). Together, our study reveals how multiple dimensions of genome evolution accompany the progressive transformation of a body plan toward the extreme of angiosperm miniaturization.

## Results

### Karyotype architecture remains broadly conserved during body plan reduction

Genomic resources within the *Wolffia* lineage have remained limited, constraining lineage-level understanding of genome evolution across duckweeds. We therefore generated a chromosome-scale, haplotype-resolved reference genome for *Wolffia globosa*. The 1.0-Gb assembly anchored 92.9% of the assembled sequence to 20 pseudomolecules with high continuity and completeness (N50 = 50.8 Mb, 93.7% BUSCO; Fig. 1a; Fig. S1; Tables S1–S5). The *W. globosa* genome contained 21,323 predicted protein-coding genes and was TE-rich (70.1%; Tables S6–S7), toward the upper end of the 15–85% range reported across plant genomes (Huang et al., 2025).

**Fig. 1.**
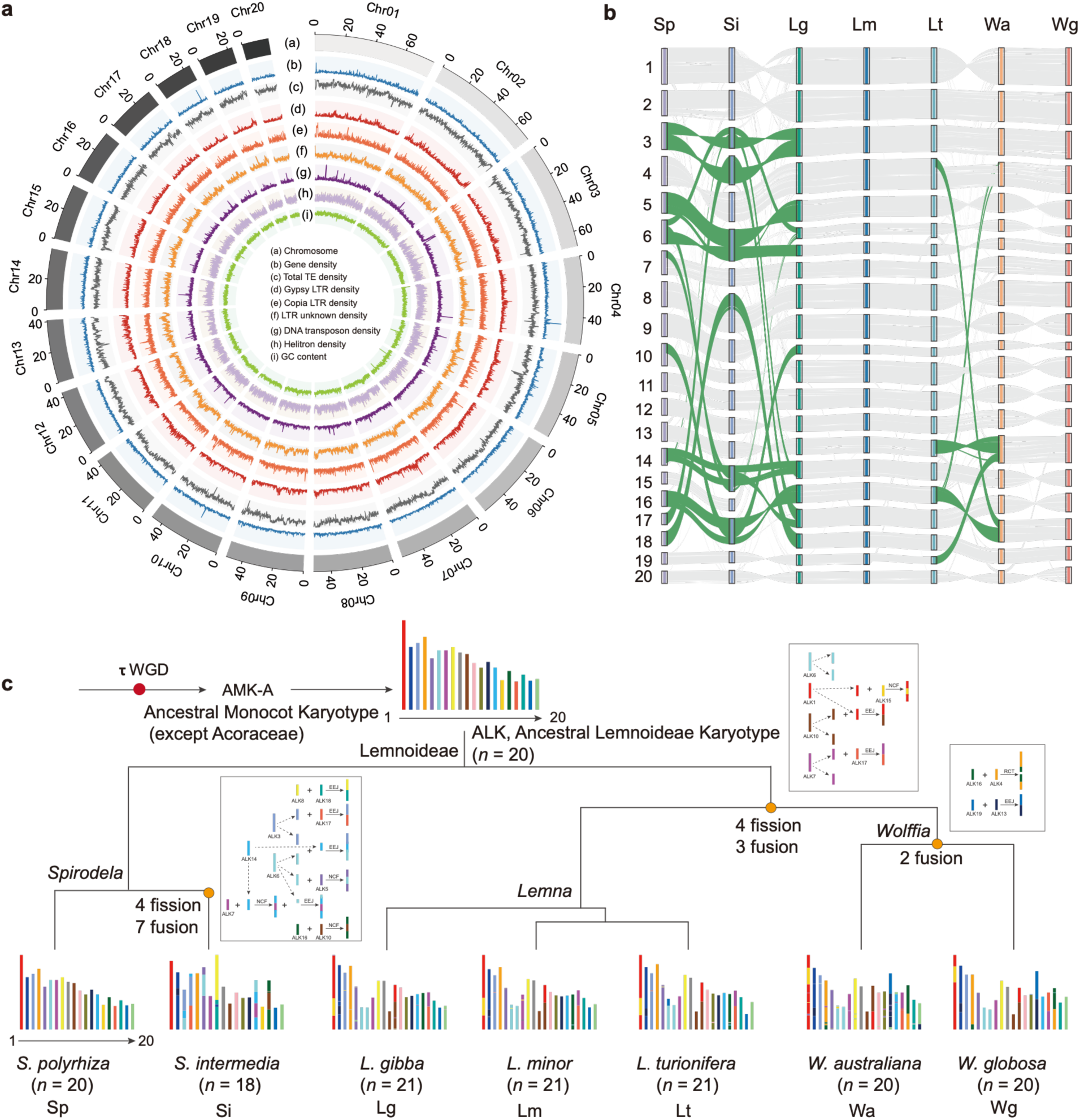
Chromosome-scale genome of *Wolffia globosa* reveals broadly conserved karyotype architecture across duckweed lineages. **a,** Chromosome-scale landscape of the *W. globosa* genome. **b,** Chromosome-level synteny across the seven duckweed species. Grey links indicate conserved synteny and green links highlight major chromosome rearrangements. **c,** Reconstruction of karyotype evolution across the evolutionary trajectory from the *Spirodela* through *Lemna* to *Wolffia* lineages. Four fissions and three fusions were reconstructed before the divergence of the *Lemna* and *Wolffia* lineages, followed by two fusions along the branch leading to the *Wolffia* lineage. Insets show inferred rearrangements, including end–end joining (EEJ), nested chromosome fusion (NCF), and reciprocal chromosome translocation (RCT). Chromosome numbers are indicated in parentheses.

Despite pronounced differences in body plan and genome size, karyotype architecture remained broadly conserved across duckweeds, showing extensive chromosome-level synteny (Fig. 1b,c; Fig. 2a). Against this conserved background, several lineage-specific fissions and fusions marked major evolutionary transitions. Four fissions and three fusions were inferred in the common ancestor of the *Lemna* and *Wolffia* lineages following divergence from the *Spirodela* lineage (Fig. 1c; Fig. 2a). Only two additional fusions were inferred in the *Wolffia* ancestor, with no subsequent fission or fusion detected within the sampled *Lemna* or *Wolffia* lineages. By contrast, the greatest karyotype reorganization occurred within the *Spirodela* lineage. Karyotype evolution thus accompanied duckweed diversification but did not progressively increase with body plan reduction toward *Wolffia*. Chromosome-scale reorganization alone therefore does not mirror the contrasting trajectories of genome expansion and progressive body plan simplification.

**Fig. 2.**
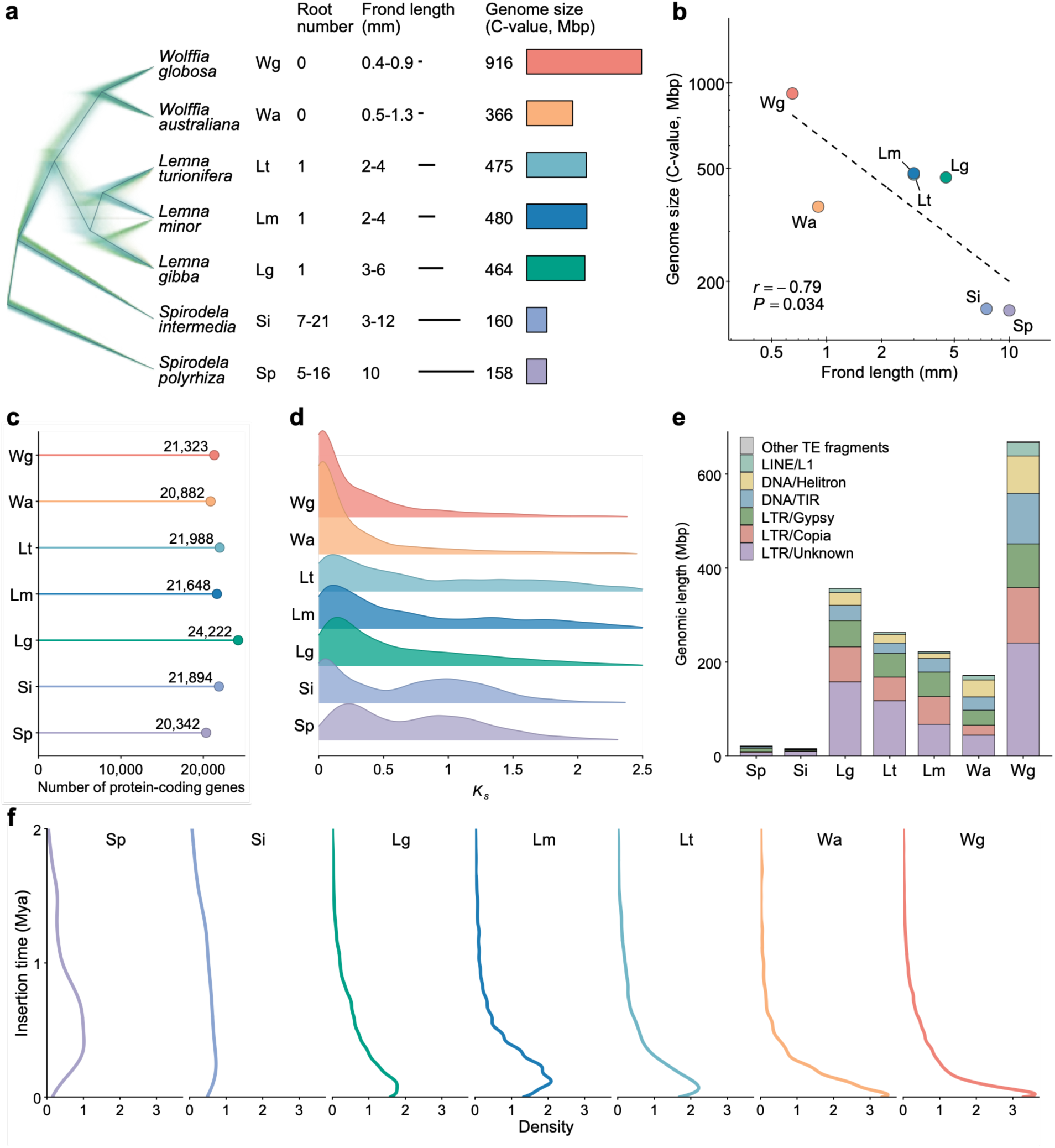
Genomic dimensions become decoupled during duckweed miniaturization. **a,** Species tree inferred from 7,365 single-copy orthologues shows progressive reduction in frond size and root number from the *Spirodela* through *Lemna* to *Wolffia* lineages, accompanied by a nearly sixfold increase in genome size. C-values represent haploid nuclear DNA content. **b,** Genome size is negatively correlated with frond length across duckweeds, revealing opposing trajectories of genome expansion and body plan reduction. **c,** Protein-coding gene numbers remain comparatively stable across lineages. **d,** Genome-wide distributions of synonymous substitution rates (*K*s) among syntenic gene pairs indicate recent gene duplication rather than recent whole-genome duplication. **e,** Transposable element (TE) abundance increases markedly with genome size, with long terminal repeat (LTR) retrotransposons accounting for much of the expanded repetitive fraction. **f,** Insertion time distributions of intact LTR retrotransposons reveal recurrent, lineage-specific amplification, with pronounced recent accumulation in the *Wolffia* lineage.

### Genome expansion reveals contrasting genomic trajectories

Progressive body plan reduction across duckweeds was accompanied by a striking increase in genome size (Fig. 2a,b). From the *Spirodela* to *Wolffia* lineages, the frond—the highly reduced vegetative body lacking conventional differentiation into stem and leaves—decreased in size by nearly an order of magnitude (Fig. 2a; Fig. S1) (Bog et al., 2020). Over the same trajectory, multiple roots were progressively reduced to complete root loss (Ware et al., 2023). Genome size moved in the opposite direction, increasing nearly sixfold. Genome size was strongly negatively correlated with frond size (Pearson’s *r* = −0.79, *P* = 0.034; Fig. 2b). This opposing pattern raised the question of what accounted for genome expansion.

Protein-coding gene number remained remarkably stable across duckweeds (20,000– 25,000; Fig. 2c) (Ernst et al., 2025) despite progressive body plan simplification, illustrating the “G-value paradox” within a single evolutionary lineage.

We next asked whether genome expansion could reflect recent whole-genome duplication (WGD). *K*s distributions showed no evidence of recent WGD (Fig. 2d). Although polyploidy is common in natural duckweed populations (Hoang et al., 2022), it represents recent population-level variation rather than a shared WGD. Instead, pronounced low *K*s peaks indicated recent gene duplication. A broader ancient duplication peak was retained in *Spirodela* but absent from *Lemna* and *Wolffia* lineages, raising the possibility of an additional WGD in the *Spirodela* lineage. However, most homologs retained 1:1 correspondence across syntenic regions both between and within lineages, rather than the 2:1 pattern expected if a *Spirodela*-specific WGD had occurred (Fig. S2). The ancient duplication signal is instead more consistent with a WGD predating duckweed diversification (Wang et al., 2014), followed by differential retention of duplicated regions among lineages. Consistent with this interpretation, duplicated synteny progressively declined from *Spirodela* through *Lemna* to *Wolffia* lineages (Fig. S2). The progressive erosion of duplicated synteny may have been facilitated by TE accumulation, as suggested by previously reported TE-associated synteny disruption in *W*. *australiana* (Michael et al., 2021).

We therefore examined whether TE accumulation could account for genome expansion. TEs increased strikingly in both relative abundance and absolute genomic content across duckweeds (Fig. 2e; Table S8). TE content rose from 12–15% in *Spirodela* to 60–70% in *Lemna* and 70% in *W. globosa*, while the absolute amount of TE-derived sequence increased with genome size. TE accumulation was dominated by LTR retrotransposons, particularly Gypsy and Copia, which are widespread across plant genomes (Huang et al., 2025; Zhang et al., 2026). The contrast within *Wolffia* was particularly striking: *W. globosa* and *W. australiana* contained similar numbers of protein-coding genes, yet the *W. globosa* genome was nearly threefold larger, with much of this difference associated with greater TE abundance. This pattern suggested more recent or extensive TE amplification in *W. globosa*. LTR insertion histories supported this possibility, revealing recurrent but progressively more recent amplification from *Spirodela* through *Lemna* to *Wolffia* (Fig. 2f). Because recently amplified elements have less time to be removed through evolutionary processes such as purifying selection (Zhang et al., 2026), recent LTR proliferation provides a plausible explanation for the exceptionally large genome of *W. globosa*. The TE-rich *W. globosa* genome was also enriched in species-specific gene families associated with telomere maintenance and DNA repair, suggesting accompanying changes in genome maintenance functions (Fig. S3). More broadly, progressive TE accumulation from *Spirodela* through *Lemna* to *Wolffia* paralleled genome expansion across duckweed lineages.

### Aquatic adaptation converges on functional simplification

Duckweed miniaturization evolved within a transition to aquatic life, raising the question of how much of its body plan simplification reflects recurrent genomic changes associated with aquatic adaptation. We assembled a comparative framework spanning duckweeds and other floating or floating-leaved aquatic plants from multiple angiosperm lineages, together with predominantly monocot terrestrial references (Fig. 3a; Table S9). Terrestrial references were selected from major monocot lineages to provide phylogenetically relevant comparisons with duckweeds and minimize differences arising from deep evolutionary divergence. *Arabidopsis thaliana* was additionally included to facilitate functional annotation and interpretation. In contrast, non-duckweed aquatic species were selected to represent floating or floating-leaved lifestyles across diverse angiosperm lineages, including monocots, eudicots and the early-diverging angiosperm *Nymphaea colorata*. This phylogenetic breadth allowed recurrent genomic changes associated with aquatic adaptation to be distinguished from lineage-specific evolution. Within Araceae, the free-floating *Pistia stratiotes* provided a close aquatic reference to distinguish duckweed-specific changes from those inherited more broadly within Araceae. CAFE analysis revealed widespread gene family expansion and contraction across the phylogeny (Fig. 3a).

**Fig. 3.**
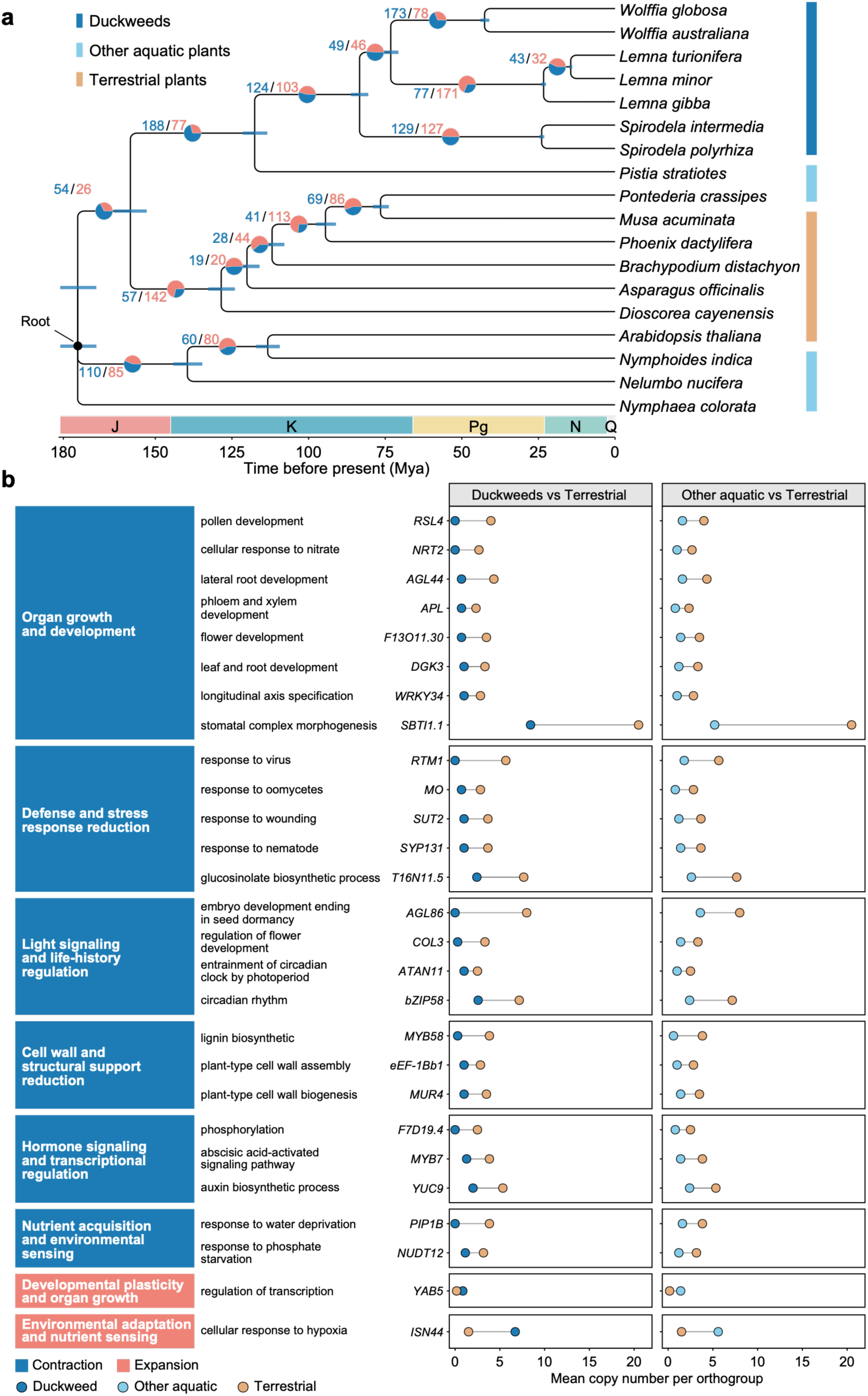
Shared functional remodeling associated with adaptation to aquatic life across duckweeds and diverse angiosperm lineages. **a,** Time-calibrated phylogenetic tree inferred from 508 single-copy orthologues places duckweeds within a nested comparison of floating and floating-leaved aquatic plants and terrestrial plants. Terrestrial species were selected primarily from major monocot lineages to minimize deep phylogenetic divergence from duckweeds, and *Arabidopsis thaliana* was additionally included to facilitate functional annotation and interpretation. Other aquatic species span monocots, eudicots, and early-diverging angiosperm lineages, providing phylogenetic breadth for identifying shared changes associated with aquatic life. Gene family contraction and expansion were reconstructed using CAFE, with blue and red numbers indicating contracted and expanded families, respectively. Bars indicate 95% highest posterior density (HPD) intervals for the estimated divergence times. Geological periods are Jurassic (J), Cretaceous (K), Paleogene (Pg), Neogene (N) and Quaternary (Q). **b,** Direct comparisons of copy numbers for gene families identified using OrthoFinder reveal shared functional changes in duckweeds and other aquatic plants relative to terrestrial plants, representing recurrent functional simplification associated with aquatic life. Dots indicate mean copy number per orthogroup.

We next asked which changes were consistently associated with aquatic lifestyles. Using gene families identified by OrthoFinder (Emms et al., 2026), we directly compared gene copy numbers between duckweeds and terrestrial plants, and independently between other aquatic plants and terrestrial plants (Fig. 3b). Despite their independent evolutionary origins, duckweeds and other aquatic plants showed similar shifts in gene repertoires relative to terrestrial plants, dominated by gene family contraction rather than expansion (Fig. 3b; Table S10). These contractions converged on functions central to the terrestrial plant body plan. Gene families associated with leaf and root development, longitudinal axis specification, vascular differentiation, cell-wall assembly and lignification were recurrently reduced, paralleling reduced investment in differentiated organs, long-distance transport, and mechanical support in aquatic environments (Koga et al., 2024; Ma et al., 2024; Olsen et al., 2016; Povilus et al., 2020). Gene families involved in stomatal complex morphogenesis were also contracted, potentially related to the specialized distribution of stomata on air-facing surfaces of floating leaves (Liu et al., 2023).

Recurrent contractions extended beyond structural development to environmental response pathways. Gene families associated with auxin and abscisic acid signaling, water deprivation, pathogen and wounding responses, and phosphate-starvation responses were reduced across aquatic lineages (Guo et al., 2025; Ma et al., 2024; Olsen et al., 2016; Tek et al., 2026). These changes point to broad remodeling of regulatory and stress-response systems that evolved in terrestrial plants for gravity-directed growth, desiccation, biotic antagonism, and root-centered nutrient acquisition. In contrast to this pervasive contraction, relatively few functions showed recurrent expansion. These included transcriptional regulation and hypoxia responses, potentially reflecting increased regulatory flexibility and adaptation to oxygen limitation in aquatic environments (Chen et al., 2026; Ma et al., 2024).

Together, these contractions define a recurrent genomic signature of aquatic adaptation, but their occurrence in structurally more complex aquatic plants indicates that they alone cannot explain the extreme reduction of the duckweed body plan.

### Duckweed miniaturization extends beyond aquatic adaptation

We next asked what genomic changes distinguished duckweed simplification from broader aquatic adaptation. Compared with other floating or floating-leaved aquatic plants, gene family contractions were more than twice as frequent as expansions in duckweeds (Fig. 4; Tables S11– S12). Contractions were concentrated in developmental and structural functions, with several functions already reduced during aquatic adaptation showing further reduction.

**Fig. 4.**
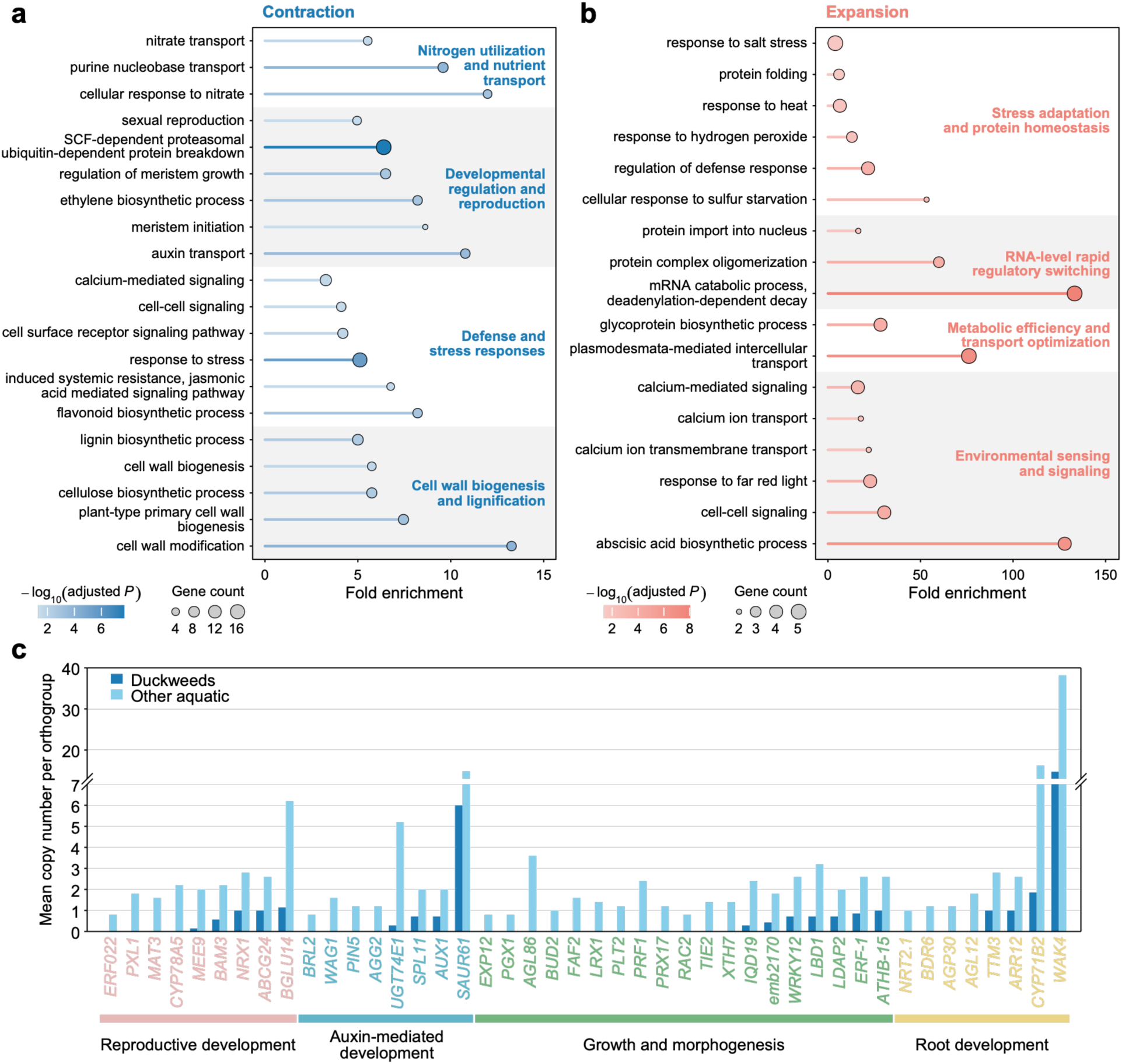
Functional streamlining in duckweeds extends beyond shared aquatic adaptation. **a,** Gene Ontology (GO) enrichment of gene families contracted in duckweeds relative to other aquatic plants reveals further functional streamlining. Fold enrichment is shown on the *x*-axis; bubble size indicates gene count and color indicates enrichment significance. **b,** GO enrichment of gene families expanded in duckweeds relative to other aquatic plants reveals selective reinforcement. **c,** Growth-and development-related gene families contributing to the functional contractions identified in **a**. Families are grouped by biological function. Bars indicate mean\ copy number per orthogroup.

Developmental contractions were particularly evident in auxin-mediated functions. Given the central roles of auxin in organ growth and vascular patterning (Barbez et al., 2012), these contractions paralleled the reduced stature and simplified vascular organization of the duckweed body plan. Root-development gene families were also further reduced as the duckweed root system progressively simplified, culminating in complete root loss in the *Wolffia* lineage.Contractions in cell wall biogenesis and lignification further accompanied the reduced vascular and structural complexity of duckweeds. Meristem-associated families were reduced, potentially reflecting a narrower repertoire of organ-initiation programs rather than diminished meristem activity (Ernst et al., 2025; Michael et al., 2021). Reproductive functions were reduced, reflecting the predominant clonal propagation of duckweeds, while contractions in nutrient utilization and transport accompanied root system simplification. Further contractions in duckweeds affected biotic defense. Gene families associated with jasmonate-mediated defense were reduced relative to other aquatic plants, in line with previously reported attenuation of canonical defense functions in duckweeds (Fang et al., 2025; Tek et al., 2026). This reduced investment in biotic defense may be associated with the exceptionally rapid growth of duckweeds, reflecting the growth–defense trade-off.

Against this pervasive contraction, a smaller set of gene families expanded in functions associated with growth, regulation, and environmental response (Fig. 4b; Table S13). Expanded families included mRNA turnover, biosynthesis, and intercellular transport, potentially supporting the rapid growth and biomass accumulation of duckweeds. Abiotic stress-response functions, including salt, heat, and sulfur stress, ABA biosynthesis, and environmental sensing, also expanded, contrasting with reduced biotic defense and potentially supporting adaptation to fluctuating aquatic conditions. Calcium-associated functions were selectively remodeled, with expansion of calcium transport and local signaling components but contraction of long-distance wound-associated signaling, suggesting enhanced environmental sensing but reduced systemic defense signaling (Gjetting et al., 2020; Shao et al., 2020). Thus, despite widespread gene family contraction, selective expansions were concentrated in functions associated with rapid growth and abiotic adaptation.

Duckweed body plan simplification therefore extended the genomic reductions shared across aquatic plants, with further contraction of developmental, structural, and biotic defense functions accompanied by selective expansion of functions associated with growth and abiotic adaptation. Whether these changes arose early in duckweed evolution or accumulated progressively toward extreme reduction in the *Wolffia* lineage remained unresolved.

### Body plan reduction accumulates through successive genomic changes

We next asked whether the genomic changes underlying duckweed miniaturization arose in a single evolutionary shift or accumulated progressively. Using CAFE-inferred gene family expansions and contractions across the phylogeny (Fig. 3a), we examined two successive transitions corresponding to major morphological reductions: the divergence of the *Lemna*– *Wolffia* lineage from the *Spirodela* lineage, followed by the divergence between the *Lemna* and *Wolffia* lineages (Fig. 5). These transitions capture progressive reductions in body size and root number. Progressive contraction could occur through two distinct modes. In the first, the same gene families contracted at both successive transitions, producing monotonic contraction. In the second, different gene families contracted at different transitions but converged on the same biological processes, producing cumulative streamlining at the process level.

**Fig. 5.**
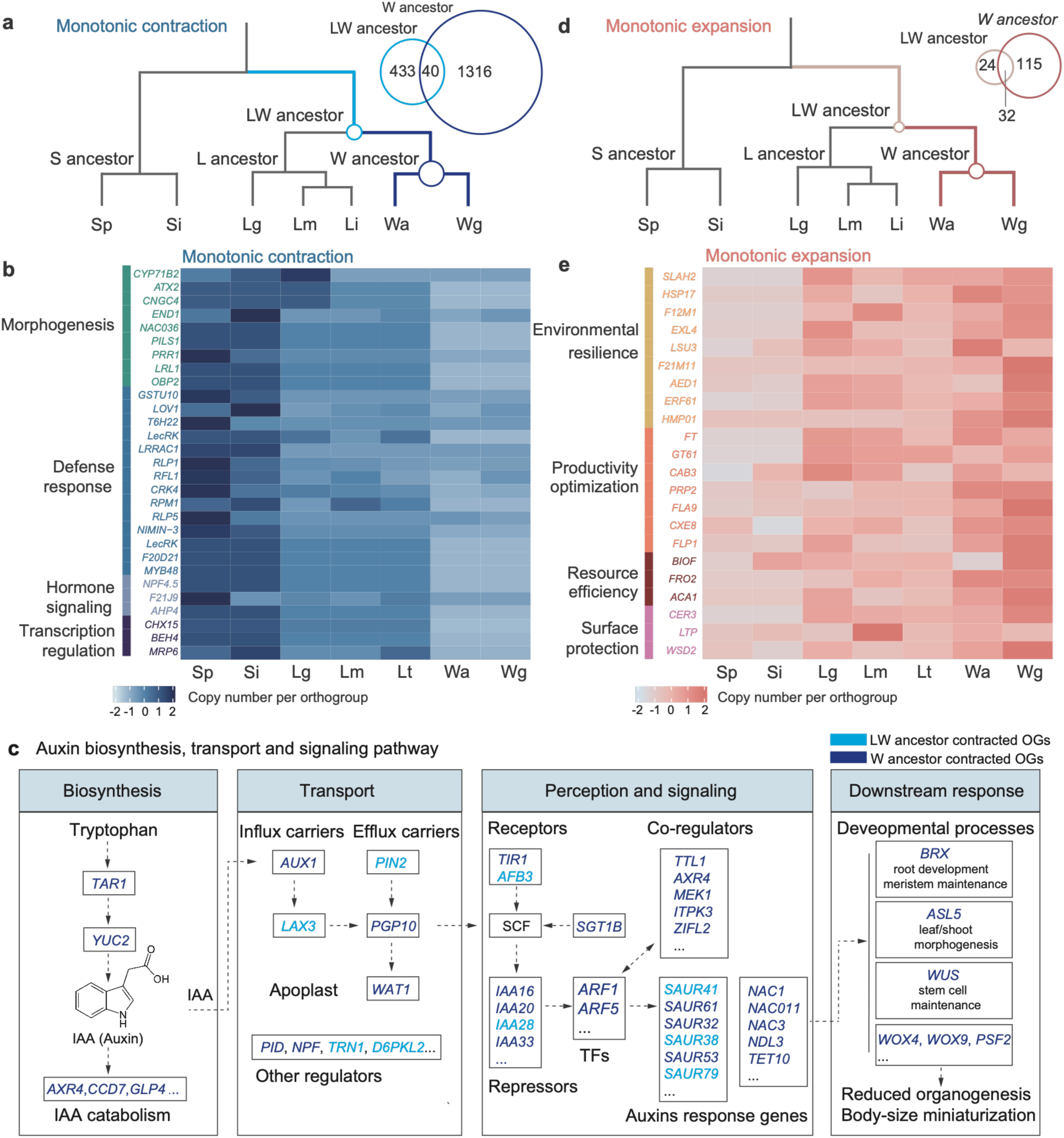
Evolutionary modes of functional streamlining during duckweed miniaturization. **a,** Gene family contractions across successive evolutionary transitions based on CAFE analysis. Light and dark blue indicate families contracted before the divergence of the *Lemna* and *Wolffia* lineages and along the branch leading to the *Wolffia* lineage, respectively. Overlapping families represent monotonic contraction across both transitions, whereas non-overlapping families represent contractions specific to either transition. Sp, *Spirodela polyrhiza*; Si, *Spirodela intermedia*; Lg, *Lemna gibba*; Lm, *Lemna minor*; Lt, *Lemna turionifera*; Wa, *Wolffia australiana*; Wg, *Wolffia globosa*. **b,** Gene families showing monotonic contraction, grouped by biological function. Heatmaps show z-score standardized copy numbers per gene family. **c,** Cumulative contraction of different gene families affecting the same biological processes, illustrated by auxin regulation. Different components of auxin biosynthesis, transport, perception and signaling, and downstream developmental regulation contracted at different transitions. Light and dark blue indicate gene families contracted at the two respective transitions. **d,** Gene family expansions across successive evolutionary transitions. **e,** Gene families showing monotonic expansion, grouped by biological function. Heatmaps show z-score standardized copy numbers per gene family.

Both modes were evident. Monotonic contraction provided the first signature of progressive genomic reduction. Gene family contraction occurred at the two transitions but became substantially more extensive toward the *Wolffia* lineage (Fig. 5a; Tables S14–S15). Forty families contracted at both transitions, representing monotonic contraction of the same gene families. These families were enriched in morphogenesis, hormone signaling, transcriptional regulation, and defense responses (Fig. 5b). Notably, this continuous reduction revealed how functions implicated in aquatic adaptation (Fig. 3b) and further reduced during duckweed simplification (Fig. 4a) were progressively remodeled across duckweed evolution. *LRL1*, involved in root hair development, and *PILS1*, involved in intracellular auxin homeostasis, contracted at both transitions, as did *BEH4*, a brassinosteroid-regulated transcription factor linked to growth and auxin signaling (Barbez et al., 2012; Galstyan & Nemhauser, 2019; Haghir et al., 2025). Thus, the same components of developmental regulation were progressively reduced across successive stages of duckweed miniaturization.

More commonly, most contracted families were specific to one of the two transitions (Fig. 5a), but collectively affected the same biological processes. Auxin regulation provided a prominent example (Fig. 5c and Fig. S4). Different components of auxin biosynthesis, transport, perception and signaling, and downstream developmental regulation contracted at different transitions, including *TAR1*, *PIN2*, *AFB3*, and *WOX4* (Chen et al., 1998; Ji et al., 2010; Swarup et al., 2008). Given the central role of auxin in organ growth and patterning, these cumulative changes paralleled the progressive reduction of roots and vascular tissues in duckweeds. Similar transition-specific contractions affected brassinosteroid biosynthesis, signaling, and downstream growth regulation (Fig. S5), suggesting cumulative remodeling of multiple hormone pathways controlling plant growth and body plan development.

Gene family expansion followed a more selective trajectory (Fig. 5d,e; Tables S14–S15). Functions expanded in duckweeds relative to other aquatic plants (Fig. 4b) were selectively reinforced across the evolutionary transitions, particularly those associated with environmental resilience and physiological performance. These included heat-shock proteins involved in stress tolerance, *SLAHs* associated with anion transport and environmental response, and functions related to resource acquisition and metabolism (Negi et al., 2008; Pinon et al., 2005; Robinson et al., 1999). Light-harvesting and surface-associated functions also expanded, including *CAB3* and *CER3*, suggesting that progressive structural simplification was accompanied by selective reinforcement of physiological functions (e.g., photosynthesis and cuticular wax biosynthesis) important to the floating lifestyle (Borisjuk et al., 2018; Kim et al., 2019; Umate, 2010).

Together, these patterns reveal two routes to progressive simplification: monotonic contraction of the same gene families and cumulative contraction of different families affecting the same biological processes. Rather than a single evolutionary shift, miniaturization emerged through stepwise genomic remodeling across duckweed evolution.

## Discussion

Our results reveal that different dimensions of genome evolution can become decoupled during organismal simplification. Across duckweed evolution, progressive reduction of the body plan was accompanied not by genome reduction, but by genome expansion driven largely by TE accumulation, while protein-coding gene number remained comparatively stable. What tracked simplification instead was the composition of the functional repertoire. Gene family contractions associated with aquatic adaptation became more pronounced in duckweeds, particularly across developmental, structural, and biotic defense functions, while selective expansions favored functions associated with growth and abiotic adaptation. These functional changes accumulated progressively through both monotonic contraction of the same gene families and cumulative contraction of different families affecting the same biological processes. Thus, genome size, gene number, and functional repertoire followed distinct evolutionary trajectories, revealing that extreme organismal miniaturization can arise through functional streamlining within an expanding genome.

Aquatic adaptation itself promotes functional simplification, but it establishes only a baseline. Duckweeds pushed this streamlining much further. Repeated transitions from terrestrial to aquatic life have been accompanied by recurrent reductions in functions associated with structural support, organ development, and terrestrial stress responses (Chen et al., 2026; Guo et al., 2025; Meseguer et al., 2022). Our comparisons across floating and floating-leaved plants reinforce this pattern. Yet these changes were shared by aquatic plants that retain far greater structural complexity than duckweeds. Relative to these aquatic plants, duckweeds underwent further contraction of developmental and structural functions, particularly auxin-mediated regulation and root development, together with reproduction and biotic defense. Importantly, this deeper streamlining was not simply a process of loss: functions associated with rapid growth and abiotic adaptation expanded selectively, suggesting a shift from investment in structural complexity and biotic defense toward growth and environmental responsiveness. Previous comparisons within duckweeds identified reductions in roots, stomata, phytohormones, and lignocellulose-associated functions (Fang et al., 2025; Michael et al., 2021; Park et al., 2021), but could not determine which changes reflected broader aquatic adaptation and which represented further remodeling associated with extreme simplification. Our broader comparison separates these evolutionary layers, showing that duckweed miniaturization built upon, but substantially exceeded, the functional streamlining associated with aquatic life.

The trajectory toward miniaturization reveals how functional streamlining can accumulate across evolutionary time. Rather than arising through a single genomic shift, streamlining accumulated across successive transitions through both monotonic contraction of the same gene families and, more commonly, contraction of different families affecting the same biological processes (Fig. 5). Thus, progressive simplification can emerge from distinct genomic changes that converge at the level of biological processes. This may be particularly consequential for highly connected regulators such as auxin and brassinosteroids, whose effects span organ initiation, growth, vascular patterning, and root development (Chen et al., 1998; Ji et al., 2010; Swarup et al., 2008), consistent with previous work implicating auxin regulation in the granule body form of *Wolffia australiana* (Park et al., 2021). Extreme miniaturization may therefore emerge not through loss of a discrete set of “body-plan genes,” but through cumulative remodeling of the developmental systems that construct the body plan.

In summary, duckweed evolution reveals how the long-standing disconnect between genomic and organismal complexity unfolds across successive stages of body plan reduction. The stronger association of organismal complexity with protein family and domain diversity (Alvarez-Ponce & Krishnamurthy, 2025) offers one explanation for the C-value and G-value paradoxes (Gregory, 2005; Hahn & Wray, 2002). Our findings add a further dimension to this view: as organismal complexity declined, the functional repertoire was not simply reduced, but selectively remodeled across successive evolutionary transitions. Organismal complexity may therefore depend not only on the size of the functional repertoire, but on how its components are reconfigured through evolution.

## Materials and Methods

### Plant material and genome assembly

An accession of *Wolffia globosa* (CH.SD.06.WO) was collected from Shandong Province, China. The axenic clonal line was established and maintained in sterile 0.5× Hoagland medium at 25 °C under a 16-h light/8-h dark photoperiod (Wei & Tan, 2023). Genome size was estimated by flow cytometry as previously described (Wei et al., 2020). Illumina short reads, PacBio HiFi reads, Hi-C data, and RNA-seq data were generated for genome characterization, *de novo* assembly, chromosome-scale scaffolding, and gene annotation. Protein-coding gene models were generated by integrating *ab initio* gene prediction, transcript evidence, and protein homology. Predicted protein-coding genes were functionally annotated by integrating sequence similarity, protein-domain, and orthology-based evidence. Details of sequencing, assembly, and annotation are provided in the Supplementary Methods.

### Duckweed phylogeny and karyotype evolution

Duckweeds are free-floating aquatic monocots comprising five genera (*Spirodela*, *Landoltia*, *Lemna*, *Wolffiella*, and *Wolffia*) and 37 species worldwide (Landolt, 1986). They include some of the smallest and fastest-growing flowering plants and exhibit a pronounced evolutionary gradient of body plan reduction. To examine how genome evolution accompanies progressive body plan reduction, we included all previously available chromosome-scale duckweed genomes together with the *W. globosa* genome generated here, spanning the three represented lineages of *Spirodela*, *Lemna*, and *Wolffia* (Table S9). The resulting seven species dataset comprised *Spirodela polyrhiza*, *Spirodela intermedia*, *Lemna gibba*, *Lemna minor*, *Lemna turionifera*, *Wolffia australiana*, and *Wolffia globosa*. A species tree was inferred from 7,365 single-copy orthologues identified by OrthoFinder v3.1.5 (Emms et al., 2026). Maximum-likelihood gene trees were reconstructed using IQ-TREE v2.4.0 (Minh et al., 2020) and used to infer the species tree under the multispecies coalescent framework with ASTRAL-IV v1.25.4.8 (Zhang et al., 2025).

To examine chromosome evolution across duckweeds, pairwise syntenic blocks were identified using MCscan implemented in JCVI v1.5.9 (Tang et al., 2024), and karyotype evolution along successive evolutionary transitions in duckweeds was reconstructed using WGDI v0.75 (Sun et al., 2022). Details of duckweed species, phylogenetic inference, synteny analysis, and karyotype reconstruction are provided in the Supplementary Methods.

### WGD and TE evolution within duckweeds

To test whether genome expansion during duckweed evolution was associated with additional whole-genome duplication (WGD), we examined duplication history across the seven genomes using complementary *K*s, self synteny and syntenic depth analyses. In parallel, to assess the contribution of transposable elements (TEs) to genome size variation during duckweed evolution, we compared TE abundance and composition across the seven genomes using the EDTA-based annotation pipeline (Ou et al., 2019). Details of WGD inference and TE analyses are provided in the Supplementary Methods.

### Species selection and phylogeny for comparative genomics

Because adaptation to aquatic life can itself involve body plan simplification, we sought to distinguish genomic changes shared with other aquatic plants from those specific to duckweeds during progressive miniaturization. We assembled a comparative dataset of 18 angiosperms comprising the seven duckweeds, five other aquatic plants, and six terrestrial species, selected based on phylogenetic relevance, ecological strategy, assembly quality, and gene annotation completeness (Fig. 3a; Table S9; see also Results). To establish the evolutionary framework for comparative genomic analyses, we reconstructed a time-calibrated phylogeny from 508 single-copy orthologues present in all 18 species. A maximum-likelihood tree was inferred using IQ-TREE and time calibrated using MCMCTree implemented in PAML v4.10.10 (Yang, 2007). Details of species selection and phylogenetic reconstruction are provided in the Supplementary Methods.

### Convergent gene family evolution in aquatic plants

To identify recurrent gene family changes associated with aquatic adaptation, we independently contrasted gene family copy numbers in two aquatic groups against the same terrestrial reference group: duckweeds versus terrestrial species, and non-duckweed aquatic species versus terrestrial species. Gene families showing concordant changes in both comparisons were then identified as convergent changes associated with aquatic adaptation. Gene families were classified as expanded or contracted only when they satisfied all three criteria: |log₂ fold change| ≥ 1, one-sided permutation *P* < 0.05, and consistent copy-number differences across species within each group. Details of effect size calculation, permutation testing and species-level consistency criteria are provided in the Supplementary Methods.

### Gene family evolution beyond shared aquatic adaptation

To identify gene family changes that distinguish duckweeds from changes shared more broadly across aquatic plants, we directly compared gene family copy numbers between duckweeds and non-duckweed aquatic species. Gene family expansions and contractions were identified using the same effect size threshold, one-sided permutation test and species-level consistency criteria described above. Details are provided in the Supplementary Methods.

### Modes of genomic evolution during duckweed miniaturization

To determine how gene family changes accumulated across successive transitions of duckweed miniaturization, gene family expansions and contractions were first reconstructed across the 18-species phylogeny using CAFE5 (Mendes et al., 2021). CAFE5 was fitted using a single global birth–death rate, and ancestral gene family sizes and copy number changes were inferred along each branch of the phylogeny. Gene families containing more than 200 copies in any species were excluded to reduce the influence of extreme copy numbers on birth–death model estimation. Gene family changes were then compared across successive evolutionary transitions within duckweeds to characterize their modes of accumulation during progressive miniaturization. Details are provided in the Supplementary Methods.

## Acknowledgements

We thank Zhe Wu, Tao Zhang, Haolin Han, and Rong Tao from the Wei Lab for assistance with duckweed collection maintenance and flow cytometry. This work was supported by the National Natural Science Foundation of China (32571740 to N.W.); Hubei Provincial Natural Science Foundation of China (2026AFA113 to N.W.); Chinese Academy of Sciences and Wuhan Botanical Garden, Chinese Academy of Sciences (E529990101 and E455990101 to N.W.); Wuhan Municipal Committee (E63E990101 to N.W.).

## Author contributions

N.W. conceptualized and designed the study. A.Z. and Z.T. conducted the data analyses and prepared the figures. N.W., A.Z. and Z.T. wrote the manuscript.

## Data availability

Publicly available genome assemblies and annotations used in this study were obtained from LEMNA.ORG (https://www.lemna.org/), including *Wolffia australiana* (Wa8730.v3), *Lemna turionifera* (Lt9434.v1), *Lemna minor* (Lm7210.v1), *Lemna gibba* (Lg7742a.v3) and *Spirodela polyrhiza* (Sp9509.v3). The genomes of *Pistia stratiotes* (CNA0036302), *Pontederia crassipes* (CNA0069368) and *Nymphoides indica* (CNA0046512) were obtained from the China National GeneBank DataBase (CNGBdb; https://db.cngb.org/). The *Nymphaea colorata* genome (GWHJJBQ00000000.1) was obtained from the Genome Warehouse (GWH; https://ngdc.cncb.ac.cn/gwh/). The remaining genomes were obtained from NCBI (https://www.ncbi.nlm.nih.gov/): *Spirodela intermedia* (GCA_902729315.2; PRJEB35634), *Nelumbo nucifera* (GCF_000365185.1; PRJNA168000), *Brachypodium distachyon* (GCF_000005505.3; PRJNA32607), *Musa acuminata* (GCA_904067035.1; PRJEB33461), *Asparagus officinalis* (GCF_001876935.1; PRJNA317340), *Dioscorea cayenensis* (GCF_009730915.1; PRJDB8681), *Phoenix dactylifera* (GCF_009389715.1; PRJNA322046) and *Arabidopsis thaliana* (GCA_978657495.1; PRJEB100887). Newly generated Illumina, PacBio HiFi, and RNA sequencing data are available at the National Genomics Data Center (BioProject PRJCA072097l, released upon publication). The chromosome-scale genome assembly and annotation of *Wolffia globosa* are available at Science Data Bank (ScienceDB; released upon publication). Other source data are provided with this paper.

## Supporting Information for

### Supplementary Methods

#### Plant material

To enable lineage-level reconstruction of genome evolution across duckweeds, we generated a chromosome-scale genome for *Wolffia globosa*, expanding genomic representation of the *Wolffia* lineage. *Wolffia globosa* is the only *Wolffia* species reported in China. An accession (CH.SD.06.WO) was collected from Shandong Province, China. The axenic clonal line was established and maintained in sterile 0.5× Hoagland medium at 25 °C under a 16-h light/8-h dark photoperiod (Wei & Tan, 2023). Plants were propagated clonally under the same condition to obtain genetically uniform material for genome and transcriptome sequencing.

Genome size was estimated by flow cytometry as previously described (Wei et al., 2020). *Spirodela polyrhiza* clone 7498, with a known 1C genome size of 158 Mb (Wang et al., 2014), was used as the internal standard. Nuclei were isolated by co-chopping young, fresh fronds of *Wolffia globosa* and the internal standard in 1 mL of WPB nuclear isolation buffer (Saint-Bio, Shanghai, China) using a sharp razor blade. The resulting nuclear suspension was filtered through a 40-μm nylon mesh to remove tissue debris, treated with RNase A (10 mg mL^-1^), and stained with propidium iodide (1 mg mL^-1^) in the dark for 30 min. Fluorescence intensity was measured on a BD Accuri C6 flow cytometer (BD Biosciences, San Jose, CA, USA). The 1C genome size of *W*. *globosa* was calculated from the ratio of its fluorescence intensity to that of the internal standard and estimated at 916 Mb.

#### Sequencing

Fresh, axenic fronds of *Wolffia globosa* were flash-frozen in liquid nitrogen and sent to BIOYIGENE (Bioyi Biotechnology Co., Ltd. Wuhan, China) for DNA and RNA extraction, library preparation, and sequencing. We generated complementary Illumina short-read, PacBio HiFi, Hi-C and RNA-seq datasets for genome characterization, *de novo* assembly, chromosome-level scaffolding, and gene annotation, respectively. For genome characterization, an Illumina paired-end library was sequenced on the NovaSeq 6000 platform to generate 150-bp paired-end reads. A 21-mer frequency distribution was generated using Jellyfish v2.3.0 (Marçais & Kingsford, 2011) and modeled with GenomeScope2 (Ranallo-Benavidez et al., 2020) to estimate genome size, heterozygosity, and repeat content. The analysis yielded an estimated haploid genome size of 758.0–759.6 Mb and a heterozygosity estimate of 0.53% (Table S1). This k-mer based estimate was lower than the 1C genome size estimated by flow cytometry (916 Mb) and the final assembled genome (∼1.0 Gb), likely reflecting underestimation of highly repetitive sequences by the fitted k-mer frequency model. For *de novo* assembly, a PacBio HiFi SMRTbell library was prepared according to the manufacturer’s instructions and sequenced on the PacBio Revio platform. Raw reads were processed using SMRT Link v25.1, and HiFi reads with predicted read quality ≥0.99 were retained. For chromosome-level scaffolding, a Hi-C library was constructed using a proximity ligation protocol and sequenced on the Illumina NovaSeq 6000 platform. For gene annotation, total RNA was extracted and sequenced on the Illumina NovaSeq 6000 platform to generate 150-bp paired-end reads. These sequencing efforts yielded 102.84 Gb of Illumina short reads (135.40× coverage), 68.19 Gb of PacBio HiFi reads (89.78× coverage), 90.78 Gb of Hi-C reads (119.51× coverage) and 11.19 Gb of RNA-seq data (Table S2).

#### Genome assembly

To generate a chromosome-scale genome and phased haplotype assemblies for *Wolffia globosa*, PacBio HiFi reads were assembled *de novo* using hifiasm v0.25.0-r726 (Cheng et al., 2021) with default parameters. The primary assembly was cleaned using the funannotate v1.8.17 (Li & Wang, 2021) to remove redundant contigs. For chromosome-scale scaffolding, Hi-C reads were aligned to the cleaned primary assembly using Juicer v1.6 (Durand et al., 2016b). Contigs were scaffolded using 3D-DNA v180922 (Dudchenko et al., 2017) with two rounds of misjoin correction (-r 2). The resulting assembly was manually reviewed and corrected using Juicebox Assembly Tools v2.20.00 (Durand et al., 2016a) and finalized using the 3D-DNA post-review pipeline. The final assembly spanned ∼1.0 Gb, with 92.9% of the assembled sequence anchored to 20 pseudochromosomes (Table S3).

The chromosome-scale assembly was validated using Hi-C contact data and assembly metrics. Hi-C reads were processed using HiC-Pro v3.1.0 (Servant et al., 2015) with default parameters. Interaction matrices were normalized using the iterative correction and eigenvector decomposition (ICE) algorithm implemented in HiC-Pro and visualized at 500-kb resolution using HiCPlotter v0.7.1 (Akdemir & Chin, 2015). The resulting contact map showed strong intrachromosomal interactions and clear separation among the 20 pseudochromosomes, supporting their chromosome-scale organization (Fig. S1). Assembly statistics, including total assembly size, contig N50, scaffold N50, and GC content, were calculated using QUAST v5.3.0 (Gurevich et al., 2013). Genome completeness was assessed using BUSCO v6.0.0 (Manni et al., 2021) in genome mode against the embryophyta_odb10 lineage dataset. The final assembly had a contig N50 of 2.3 Mb, a scaffold N50 of 50.8 Mb, and 93.7% complete BUSCOs (Table S3).

To compare genome structure between haplotypes, the two phased assemblies generated by hifiasm were independently scaffolded against the chromosome-scale primary assembly using RagTag v2.1.0 (Alonge et al., 2022). The resulting chromosome-scale haplotype assemblies were aligned against each other using minimap2 v2.28 (Li, 2018) with the asm5 preset, and the alignments were sorted and indexed using SAMtools v1.20 (Li et al., 2009). Syntenic regions and structural variants, including inversions, translocations, duplications, insertions and deletions, were identified using SyRI v1.7 (Goel et al., 2019) with default parameters and visualized using plotsr v1.1.1 (Goel & Schneeberger, 2022).

#### Genome annotation

Repetitive sequences were first annotated to generate a repeat-masked genome for gene prediction. EDTA v2.2 (Ou et al., 2019) was used to construct a species-specific TE library by integrating structure-based and homology-based approaches. The resulting TE library was used with RepeatMasker v4.2.3 (Tarailo-Graovac & Chen, 2009) to annotate repetitive sequences and generate a soft-masked genome for subsequent gene prediction. Low-complexity sequences and non-coding RNAs were excluded from masking.

Protein-coding gene models were generated by integrating *ab initio* gene prediction, transcript evidence, and protein homology. *Ab initio* gene predictions were generated using AUGUSTUS and GeneMark-ETP implemented in BRAKER3 (Gabriel et al., 2024). RNA-seq reads were aligned to the soft-masked genome using HISAT2 v2.2.1 (Kim et al., 2019), and the alignments were sorted and indexed using SAMtools. Transcript models were reconstructed from the RNA-seq alignments using StringTie v3.0.0 (Shinder et al., 2026), and coding regions were predicted using TransDecoder v5.5.0 (Haas et al., 2013). For protein-homology evidence, protein sequences from seven previously published duckweed genomes (Table S9) were aligned to the *W*. *globosa* genome using miniprot v0.18-r281 (Li, 2023). The three sources of evidence were integrated using EVidenceModeler (EVM) v2.1.0 (Haas et al., 2008) to generate consensus gene models. These models were further refined using the PASA v2.5.3 annotation comparison pipeline (Haas et al., 2003) based on transcript alignments to improve exon–intron boundaries and gene structures. The completeness of the predicted protein-coding gene set was assessed using BUSCO v6.0.0 in protein mode against the embryophyta_odb10 lineage dataset.

Predicted protein-coding genes were functionally annotated by integrating sequence similarity, protein-domain, and orthology-based evidence. Predicted proteins were searched against the UniProtKB/Swiss-Prot and NCBI RefSeq Plant databases using DIAMOND v2.1.24 (Buchfink et al., 2015) in BLASTP mode, with an E-value cutoff of 1 × 10^-^⁵. Conserved protein domains and functional signatures were identified using InterProScan v5.59-91.0 (Jones et al., 2014), and associated Gene Ontology (GO) terms (Gene Ontology Consortium, 2021) were assigned. Orthology-based functional annotations were obtained using eggNOG-mapper v2.1.13 (Cantalapiedra et al., 2021). Functional annotation coverage across RefSeq Plant, eggNOG, InterPro, Swiss-Prot and GO is summarized in Table S6.

Non-coding RNAs were annotated separately. Transfer RNA (tRNA) genes were predicted using tRNAscan-SE v2.0.12 (Chan et al., 2021) in eukaryotic search mode. Other non-coding RNAs, including ribosomal RNAs, small nuclear RNAs, small nucleolar RNAs, microRNAs, and other conserved RNA families, were identified by searching the Rfam covariance model database using Infernal v1.1.5 (Nawrocki & Eddy, 2013) with default parameters. In total, 14,918 non-coding RNAs were annotated in the *Wolffia globosa* genome (Table S5).

#### Duckweed species and phylogeny

Duckweeds (Araceae, subfamily Lemnoideae) are free-floating aquatic monocots comprising five genera (*Spirodela*, *Landoltia*, *Lemna*, *Wolffiella*, and *Wolffia*) and 37 species worldwide (Landolt, 1986). They include some of the smallest and fastest-growing flowering plants and exhibit a pronounced evolutionary gradient of body plan reduction, from the larger, multi-rooted *Spirodela* through the reduced *Lemna* to the highly simplified *Wolffia*. To examine how genome evolution accompanies progressive body plan reduction, we included all previously available chromosome-scale duckweed genomes together with the *W. globosa* genome generated here, collectively spanning the three major evolutionary lineages represented by *Spirodela*, *Lemna*, and *Wolffia* (Table S9). The resulting seven species dataset comprised *Spirodela polyrhiza*, *Spirodela intermedia*, *Lemna gibba*, *Lemna minor*, *Lemna turionifera*, *Wolffia australiana*, and *Wolffia globosa*.

Frond length, root number, and 1C genome size were compiled for each of the seven duckweed species. Frond length was used as a proxy for body size, and root number was used to characterize progressive root reduction across the *Spirodela*, *Lemna*, and *Wolffia* lineages. These phenotypic data were compiled from Wayne’s Word (https://www.waynesword.net), the Flora of Australia (https://profiles.ala.org.au/opus/foa/), and published taxonomic descriptions (Bog et al., 2020). For genome size, the 1C value for *W*. *globosa* was determined by flow cytometry as described above, and estimates for the remaining six species were obtained from published data (Hoang et al., 2022).

To reconstruct the evolutionary history of the duckweed lineages, we inferred a species tree for the seven species under the multispecies coalescent framework. OrthoFinder v3.1.5 (Emms et al., 2026) identified 7,365 single-copy orthologues present in all seven species. For each orthologue, protein sequences were aligned using MAFFT v7.525 (Katoh et al., 2002) and the best-fitting substitution model was selected using ModelFinder. Maximum-likelihood gene trees were then built using IQ-TREE v2.4.0 (Minh et al., 2020) with 1,000 ultrafast bootstrap replicates. The resulting 7,365 gene trees were used to infer the species tree using ASTRAL-IV v1.25.4.8 (Zhang et al., 2025). To visualize gene-tree discordance relative to the inferred species tree, internal branches with bootstrap support <30% were collapsed in individual gene trees after species tree inference. The collapsed gene trees were overlaid using the R package ape (Paradis & Schliep, 2019), with the ASTRAL species tree superimposed as a reference.

#### Karyotype evolution within duckweeds

To examine chromosome evolution across duckweeds, we first compared chromosome-level synteny across the seven duckweed genomes to characterize the conservation and rearrangement of chromosome structure. Pairwise syntenic blocks were identified using MCscan implemented in JCVI v1.5.9 (Tang et al., 2024) based on protein sequence similarity and gene order. The *Spirodela polyrhiza* genome was used as a common reference to facilitate comparisons across species, given the early-diverging position of the *Spirodela* lineage within duckweeds (Tippery et al., 2015). Chromosome synteny and structural rearrangements across the seven species were visualized using the JCVI karyotype module.

We next reconstructed karyotype evolution to trace changes in chromosome organization along successive evolutionary transitions in duckweeds using WGDI v0.75 (Sun et al., 2022). Protein homology searches were performed using DIAMOND with an E-value cutoff of 1 × 10^-^⁵, and collinear blocks were identified using WGDI. The *S. polyrhiza* karyotype was used as a proxy for the ancestral Lemnoideae karyotype (ALK), based on the early-diverging position of *Spirodela* (Tippery et al., 2015). Syntenic correspondences between the ALK and the other six genomes were used to trace the conservation and reorganization of chromosome segments and to infer chromosome fissions, fusions, and translocations. Rearrangement mechanisms were classified as nested chromosome fusion (NCF), end-to-end joining (EEJ), or reciprocal chromosome translocation (RCT) according to the organization of ancestral chromosome segments in descendant chromosomes.

#### WGD inference within duckweeds

To test whether genome expansion during duckweed evolution was associated with additional whole-genome duplication (WGD), we examined duplication history across the seven genomes using complementary *K*s, self synteny, and syntenic depth analyses. Intragenomic collinear blocks were identified for each genome using WGDI, and synonymous substitution rates (*K*s) were calculated for syntenic gene pairs within collinear blocks. The distributions of *K*s values for syntenic paralogues were compared across species to identify signatures and the relative timing of large-scale duplication events. Genome-wide self synteny dot plots were generated for each species to examine the distribution and extent of duplicated chromosome segments, with syntenic gene pairs colored by their *K*s values. We further analyzed syntenic depth to distinguish lineage-specific duplication from differential retention of more ancient duplicated regions. Three representative genome pairs were compared using JCVI: *Spirodela polyrhiza*–*Lemna gibba*, *Spirodela polyrhiza*–*Wolffia australiana*, and *W. australiana*–*Wolffia globosa*. For each comparison, syntenic depth was quantified as the number of collinear genomic regions in one genome corresponding to a region in the other. A 1:1 relationship indicated conserved syntenic depth, whereas a 1:2 relationship indicated duplication.

#### TE evolution within duckweeds

To assess the contribution of transposable elements (TEs) to genome size variation during duckweed evolution, we compared TE abundance and composition across the seven genomes using the EDTA-based annotation pipeline described above. To reconstruct the temporal dynamics of LTR retrotransposon accumulation, intact LTR retrotransposons were identified using LTR_retriever (Ou & Jiang, 2018) implemented in EDTA, and their insertion times were estimated from sequence divergence between paired 5′ and 3′ LTRs. Insertion times were estimated as *T* = *K*/(2*r*), where *K* is the Kimura two-parameter distance between the paired LTRs of each intact element and *r* is the substitution rate used for dating. We used a rate of 3.04 × 10⁻⁸ substitutions per site per year (Zhang et al., 2026), derived from the experimentally estimated mutation rate of 2.38 × 10⁻¹⁰ substitutions per site per asexual generation in duckweeds (Xu et al., 2019).

#### Species selection and phylogeny for comparative genomics

Because adaptation to aquatic life can itself involve body plan simplification, we sought to distinguish genomic changes shared with other aquatic plants from those specific to duckweeds during progressive miniaturization. We assembled a comparative dataset of 18 angiosperms comprising the seven duckweeds, five other aquatic plants, and six terrestrial species, with sufficient assembly quality and gene annotation completeness (Table S9). The terrestrial species were selected predominantly from major monocot lineages to provide phylogenetically relevant comparisons with duckweeds and reduce differences attributable to deep evolutionary divergence. These species included *Musa acuminata*, *Phoenix dactylifera*, *Brachypodium distachyon*, *Asparagus officinalis*, and *Dioscorea cayenensis*. *Arabidopsis thaliana* was additionally included as a well-annotated reference to facilitate functional annotation and interpretation. Five non-duckweed aquatic species were selected to represent floating or floating-leaved lifestyles that evolved across diverse angiosperm lineages: *Pistia stratiotes*, *Pontederia crassipes*, *Nymphoides indica*, *Nymphaea colorata*, and *Nelumbo nucifera*. These species spanned monocots, eudicots and the early-diverging angiosperm lineage represented by *N. colorata*, providing broad phylogenetic breadth to identify recurrent genomic changes associated with aquatic adaptation rather than those confined to individual lineages. Within Araceae, the free-floating *P. stratiotes* provided a close aquatic reference for distinguishing duckweed-specific changes from those shared more broadly within the family.

We next reconstructed a time-calibrated phylogeny to establish the evolutionary framework for comparative genomic analyses across the 18 species. For each species, only the longest protein-coding transcript per gene was retained. OrthoFinder identified 508 single-copy orthologues present in all 18 species. Coding sequences for each orthologue were aligned separately and then concatenated. A maximum-likelihood tree was inferred using IQ-TREE. The resulting topology was fixed for divergence time estimation using MCMCTree implemented in PAML v4.10.10 (Yang, 2007). Two secondary calibration bounds from TimeTree 5 (https://timetree.org/) were used: 82–117 Ma for the divergence between *Pistia stratiotes* and the duckweed lineage, and 168–191 Ma for the divergence between *Nymphoides indica* and *Nelumbo nucifera*.

#### Convergent gene family evolution in aquatic plants

To identify recurrent gene family changes associated with aquatic adaptation, we independently contrasted gene family copy numbers in two aquatic groups against the same terrestrial reference group: duckweeds versus terrestrial species, and non-duckweed aquatic species versus terrestrial species. Gene families showing concordant changes in both comparisons were then identified as convergent aquatic-associated changes.

For each gene family in each comparison, effect size was calculated as the log₂ fold change in mean copy number between groups. A pseudocount of 0.5 was added to both group means to avoid undefined values when the mean copy number in either group was zero. A minimum effect size threshold of |log₂ fold change| ≥ 1 was applied to retain gene families showing at least a twofold difference in mean copy number between groups. Statistical support for gene family expansion or contraction was then evaluated by exhaustively permuting species labels between the two groups while preserving the original group sizes. For each gene family, the observed difference in mean copy number was compared with the permutation distribution of differences obtained from all possible label reassignments, and a one-sided permutation *P* value was calculated.

Because differences between group means may be driven disproportionately by extreme copy numbers in a subset of species rather than reflect a consistent group-level pattern, we additionally required the inferred change to be broadly supported across species within each group. For each gene family, the midpoint between the two observed group means was used as a reference to assess species-level support for the inferred change. For an inferred expansion, all but one species in the target group were required to have copy numbers above the midpoint, whereas all but one species in the reference group were required to have copy numbers below it; the criteria were reversed for an inferred contraction.

Gene families were classified as expanded or contracted only when they satisfied all three criteria: |log₂ fold change| ≥1, one-sided permutation *P* < 0.05, and the corresponding species-level consistency criterion. Repeated aquatic-associated expansions and contractions were defined as gene families that independently met these criteria in both duckweed–terrestrial and other aquatic–terrestrial comparisons. Shared gene families were annotated using *Arabidopsis thaliana* homologs identified by OrthoFinder.

#### Gene family evolution beyond shared aquatic adaptation

To identify gene family changes that distinguish duckweeds from the broader patterns shared among aquatic plants, we directly compared gene family copy numbers between duckweeds and non-duckweed aquatic species. Gene family expansions and contractions were identified using the same effect size threshold, one-sided permutation test, and species-level consistency criteria described above.

#### Modes of genomic evolution during duckweed miniaturization

To determine how gene family changes accumulated across successive transitions of duckweed miniaturization, gene family expansions and contractions were first reconstructed across the 18-species phylogeny using CAFE5 (Mendes et al., 2021). CAFE5 was fitted using a single global birth–death rate, and ancestral gene family sizes and copy number changes were inferred along each branch of the phylogeny. Gene families containing more than 200 copies in any species were excluded to reduce the influence of extreme copy numbers on birth–death model estimation.

We focused on two successive transitions corresponding to major stages of morphological reduction within duckweeds: the branch leading to the common ancestor of the *Lemna* and *Wolffia* lineages following divergence from the *Spirodela* lineage (LW transition), and the branch leading to the common ancestor of the *Wolffia* lineage following divergence from the *Lemna* lineage (W transition). Gene family contractions across these transitions were examined for two modes of progressive change. Monotonic contraction was defined as contraction of the same gene family at both the LW and W transitions. Cumulative contraction was assessed by identifying different gene families that contracted at the two transitions but affected the same biological process or developmental pathway. Families contracting at only one of the two transitions were classified as transition-specific contractions. Gene family expansions were examined across the same two transitions to assess how expanded functions accumulated across successive stages of duckweed miniaturization.

**Fig. S1.**
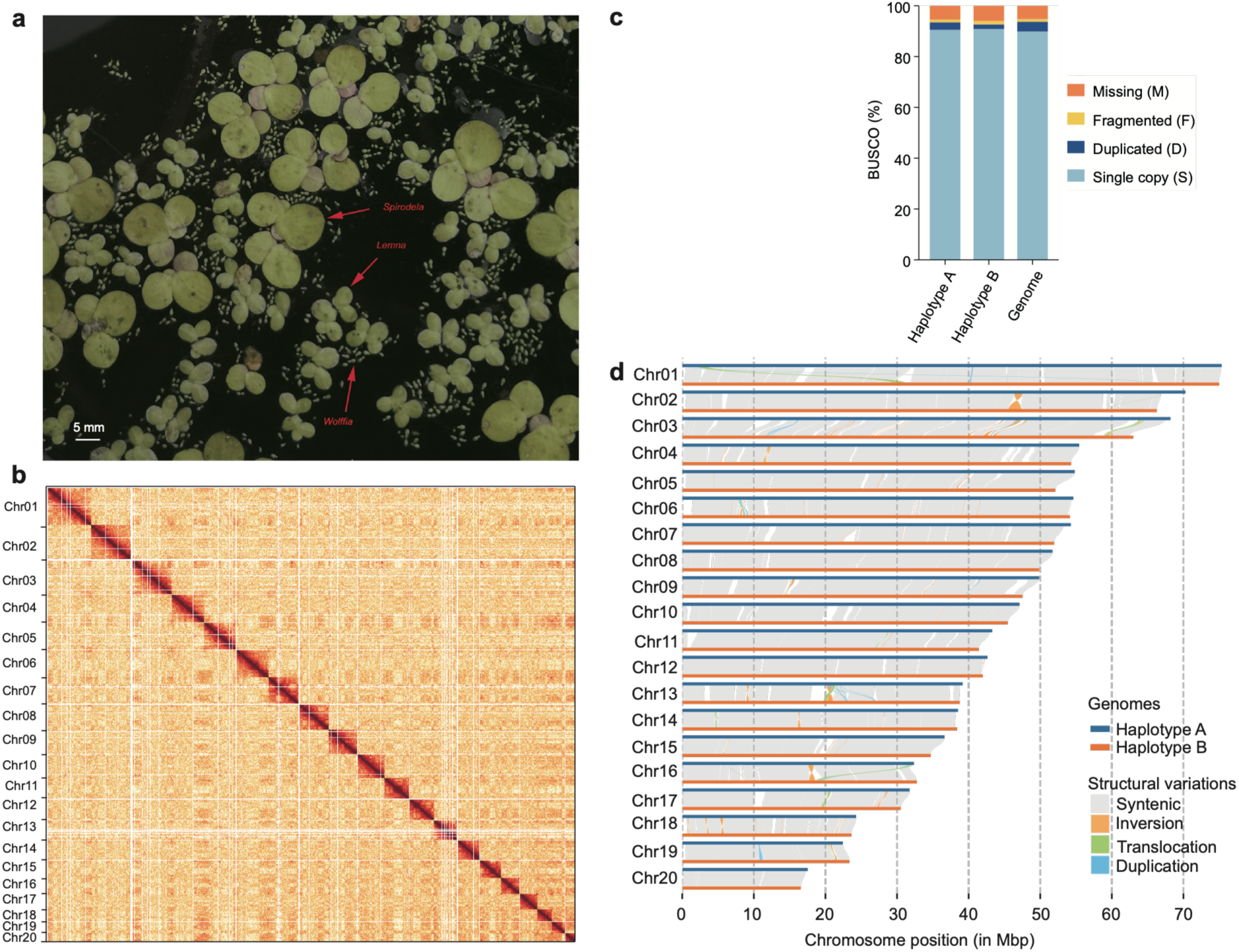
Morphological reduction across duckweed lineages and chromosome-scale genome assembly of *Wolffia globosa*. (a) Representative morphology of *Spirodela*, *Lemna* and *Wolffia* co-occurring in the same habitat, showing differences in frond size among the three lineages. (b) Hi-C contact map of the *Wolffia globosa* genome assembly at 500-kb resolution. Darker colors indicate stronger chromatin interactions. (c) BUSCO assessment of the two haplotype assemblies and the final *Wolffia globosa* genome using the embryophyta_odb10 dataset. S, complete single-copy; D, complete duplicated; F, fragmented; M, missing BUSCOs. (d) Structural variation between the two phased haplotype assemblies of *Wolffia globosa*. Gray links indicate conserved syntenic regions, whereas colored links indicate structural variants, including inversions, translocations and duplications.

**Fig. S2.**
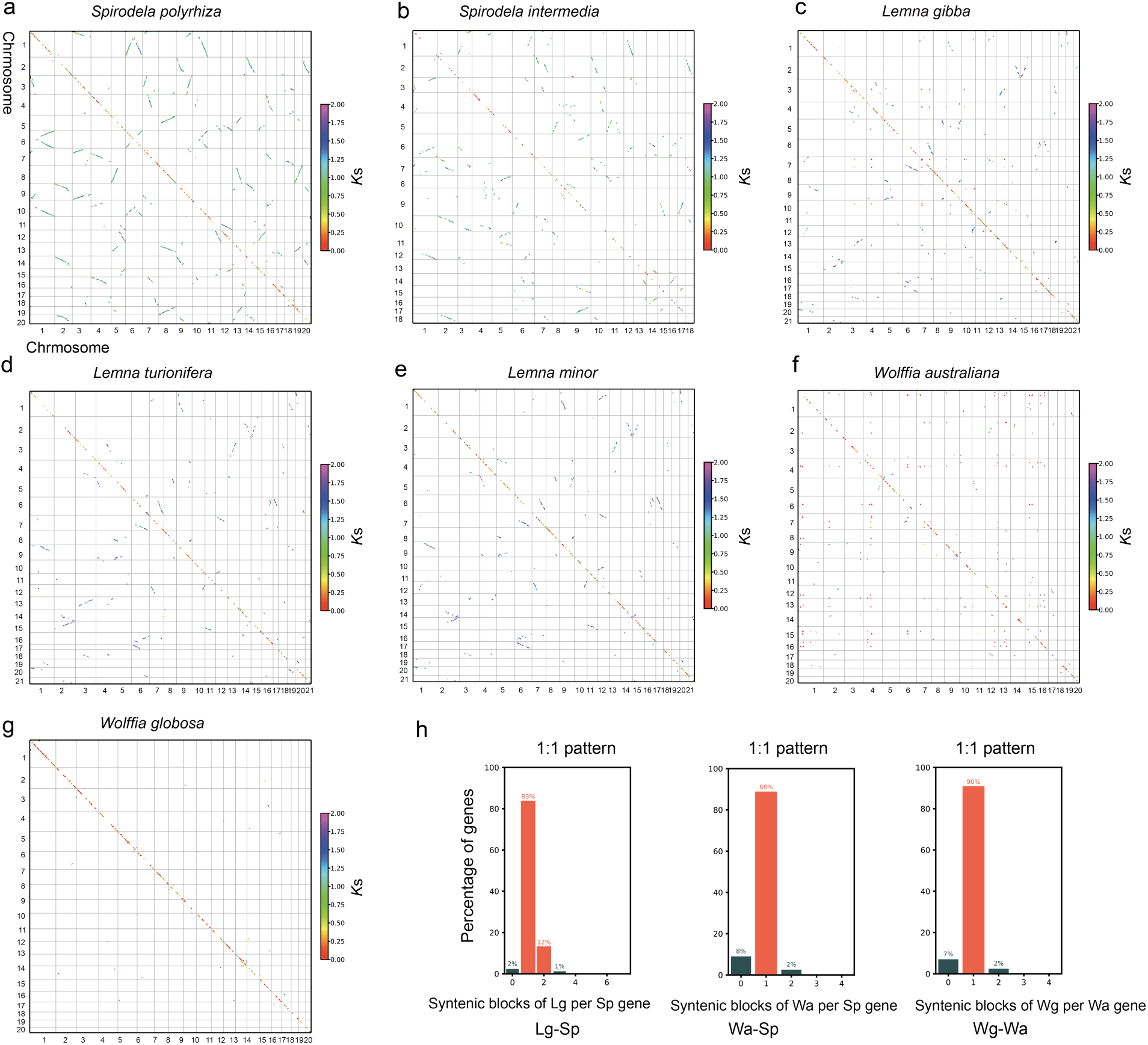
Genome-wide self synteny and syntenic depth across duckweed genomes. (a-g) Genome-wide self synteny of *Spirodela polyrhiza* (Sp)*, Spirodela intermedia*, *Lemna gibba* (Lg), *Lemna turionifera*, *Lemna minor*, *Wolffia australiana* (Wa), and *Wolffia globosa* (Wg). Each dot represents a pair of syntenic genes, colored by synonymous substitution rate (*K*s). Off-diagonal duplicated blocks progressively decrease from *Spirodela* through *Lemna* to *Wolffia*, indicating reduced retention of ancient duplicated synteny. (h) Syntenic depth between representative duckweed genomes. Bar plots show the percentage of genes at each syntenic depth in each comparison. The predominance of a syntenic depth of one supports overall 1:1 syntenic relationships in the Lg–Sp, Wa–Sp and Wg–Wa comparisons.

**Fig. S3.**
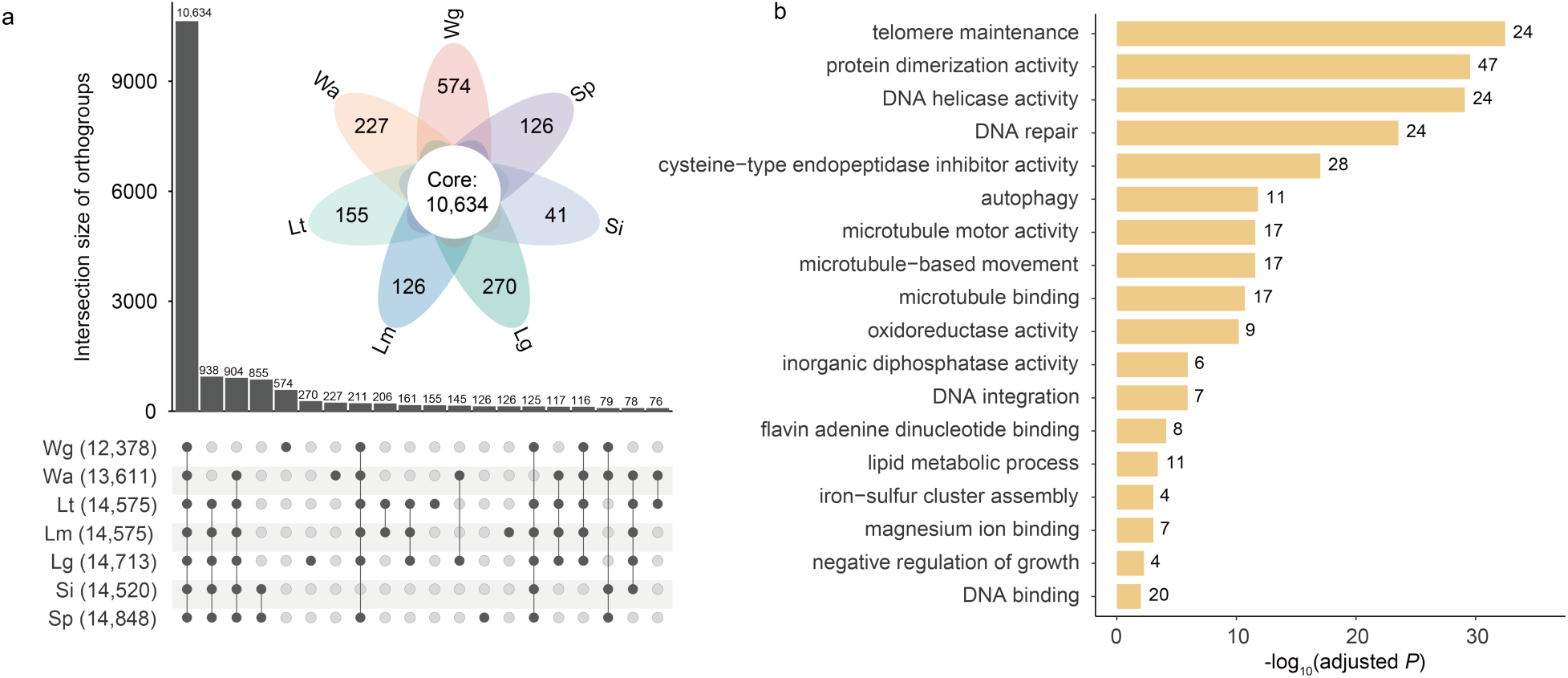
Species-specific gene families across duckweed species and functional enrichment of *Wolffia globosa*-specific genes. (a) UpSet plot showing the distribution of shared and species-specific gene families across the seven duckweed species. Numbers in parentheses indicate the total number of gene families in each species. (b) Gene Ontology (GO) enrichment of genes assigned to *Wolffia globosa* specific gene families. All GO-annotated *Wolffia globosa* protein-coding genes were used as the background. GO enrichment was assessed using a hypergeometric test with Benjamini–Hochberg correction (adjusted *P* < 0.05). Bar length represents −log_10_ (adjusted *P*), and numbers indicate the number of genes associated with each GO term.

**Fig. S4.**
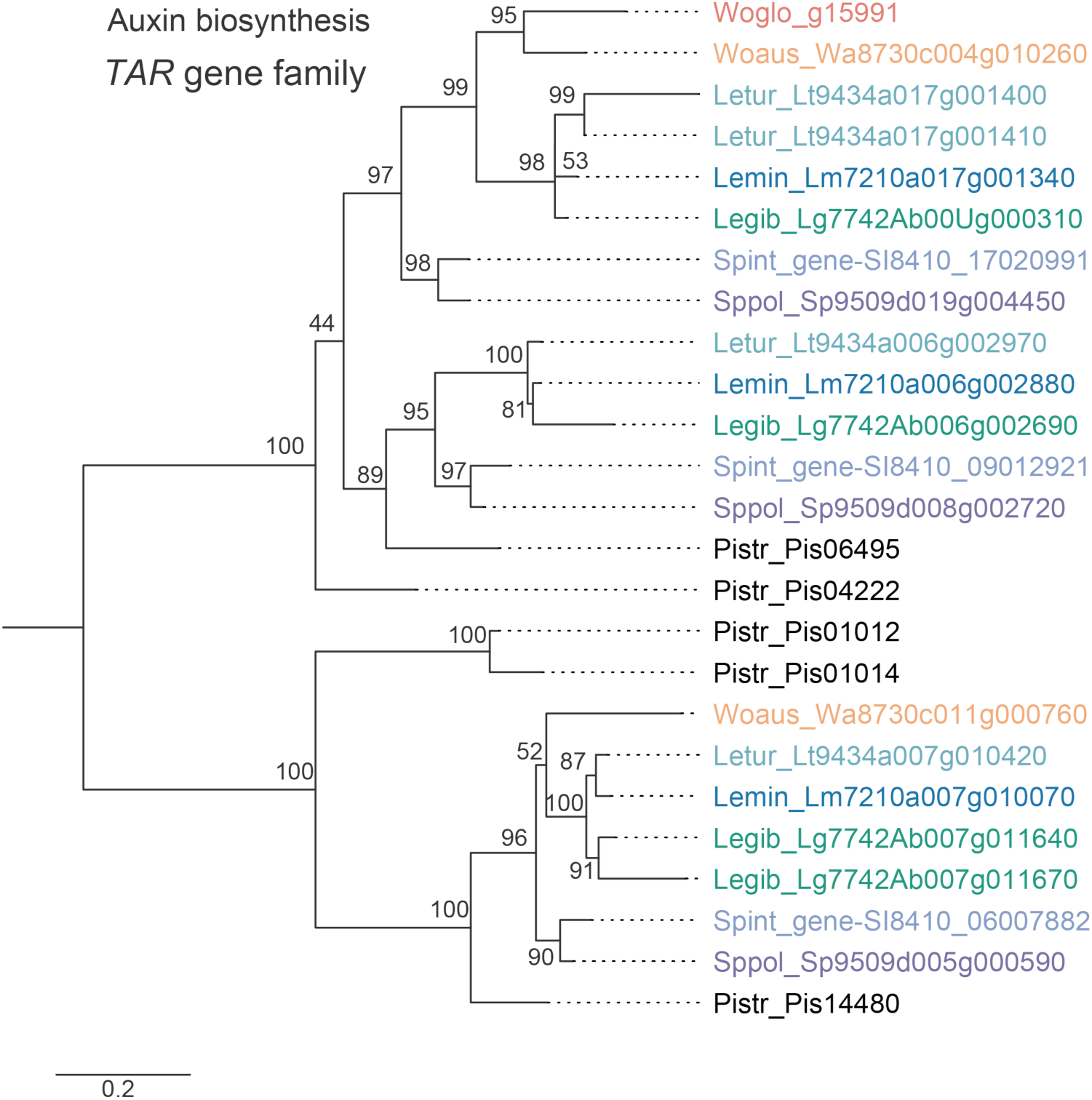
Phylogenetic analysis of the *TAR* gene family in duckweeds. Maximum-likelihood phylogeny of tryptophan aminotransferase-related (TAR) proteins from the seven duckweed species, with *Pistia stratiotes* homologs included as outgroups. Numbers at internal nodes indicate ultrafast bootstrap support values, and the scale bar represents amino acid substitutions per site. Gene labels are colored by species.

**Fig. S5.**
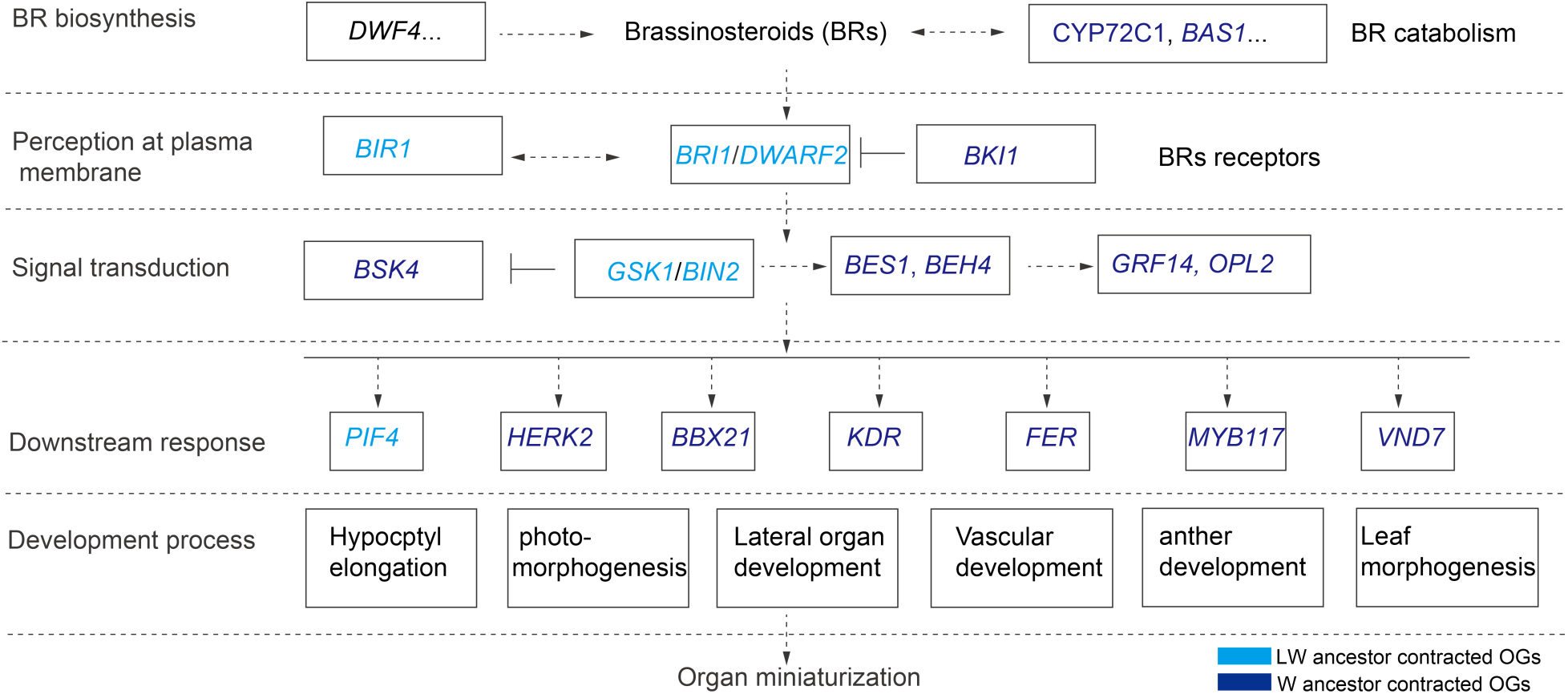
Progressive contraction across the brassinosteroid biosynthesis (BR) metabolism and signaling pathway during duckweed evolution. Gene families involved in BR biosynthesis, perception, signal transduction, downstream responses and developmental processes contracted at successive evolutionary transitions. Light and dark blue indicate families contracted at the common ancestor of *Lemna* and *Wolffia* (LW ancestor) and the common ancestor of *Wolffia* (W ancestor), respectively.

**Table S1.** Genome characterization of *Wolffia globosa* genome.

| **Property** | **Min** | **Max** |
| --- | --- | --- |
| Estimated heterozygosity rate | 0.53% | 0.56% |
| Estimated haploid genome length | 757,951,202 bp | 759,554,969 bp |
| Estimated repeat length | 278,272,524 bp | 278,861,327 bp |
| Estimated unique sequence length | 479,678,678 bp | 480,693,642 bp |
| Model Fit | 68.72% | 89.01% |

Note: Heterozygosity rate was estimated from the proportion of heterozygous (ab) k-mers relative to homozygous (aa) k-mers using the GenomeScope2 diploid model. Genome size estimates represent the haploid genome length inferred from the 21-mer frequency distribution.

**Table S2.** Sequencing data for the *Wolffia globosa* genome assembly.

| **Library** | **Usage** | **Clean Data (Gb)** | **Depth** |
| --- | --- | --- | --- |
| Illumina short reads | Genome characterization | 102,84 | 135.40× |
| PacBio HiFi | Genome assembly | 68.19 | 89.78× |
| Hi-C | Genome scaffolding | 90.78 | 119.51 |
| RNA-seq | Gene annotation | 11.19 | \ |

**Table S3.** Summary statistics of the *Wolffia globosa* genome assembly.

| **Feature** | **Genome** | **HapA** | **HapB** |
| --- | --- | --- | --- |
| Genome size (Mb) | 1,001.65 | 1,008.77 | 895.17 |
| Number of psedo-chromosomes | 20 | 20 | 20 |
| Anchored to chromosome (%) | 92.85 | 90.24% | 98.50% |
| Contig N50 (Mb) | 2.27 | 1.66 | 1.83 |
| Number of contigs | 2,770 | 2,917 | 1,013 |
| Scaffold N50 (Mb) | 50.76 | 49.87 | 49.87 |
| Number of scaffolds | 1,434 | 2,028 | 172 |
| GC content (%) | 42.8 | 42.57 | 43.12 |
| BUSCO completeness (%) | 93.7 | 93.5 | 92.7 |
| BUSCO completeness (%) of chromosomes | 93.6 | 93.3 | 92.3 |
| Genes | 21,323 | 2,0987 | 20,467 |

**Table S4.** Structural variation between haplotypes A and B of the *Wolffia globosa* genome assembly.

| **Variation_type** | **Count** | **Length_ref (bp)** | **Length_qry (bp)** |
| --- | --- | --- | --- |
| Syntenic regions | 274 | 832,747,619 | 813,053,784 |
| Inversions | 128 | 8,082,346 | 8,369,797 |
| Translocations | 138 | 6,129,749 | 6,357,668 |
| Duplications of HapA | 61 | 380,642 | - |
| Duplications of HapB | 279 | - | 2,270,854 |
| Not aligned of HapA | 549 | 64,626,982 | - |
| Not aligned of HapB | 746 | - | 52,133,532 |

Sequence variation
| **Variation_type** | **Count** | **Length_ref (bp)** | **Length_qry (bp)** |
| --- | --- | --- | --- |
| SNPs | 1,202,846 | 1,202,846 | 1,202,846 |
| Insertions | 72,400 | - | 9,846,993 |
| Deletions | 73,582 | 13,152,051 | - |
| Copygains | 43 | - | 375,004 |
| Copylosses | 44 | 1,212,751 | - |
| Highly diverged | 3,814 | 131,097,869 | 115,924,999 |
| Tandem repeats | 11 | 52,638 | 39,187 |

**Table S5.** Summary of non-coding RNA identified in the *Wolffia globosa* genome assembly.

| **Class** | **Subclass** | **Number** |
| --- | --- | --- |
| tRNA |  | 10,575 |
| rRNA |  |  |
|  | 5S_rRNA | 1,135 |
|  | 5_8S_rRNA | 244 |
|  | LSU_rRNA_eukarya | 1,221 |
|  | SSU_rRNA_eukarya | 1,161 |
| miRNA |  | 72 |
| snoRNA |  | 2,490 |

**Table S6.**
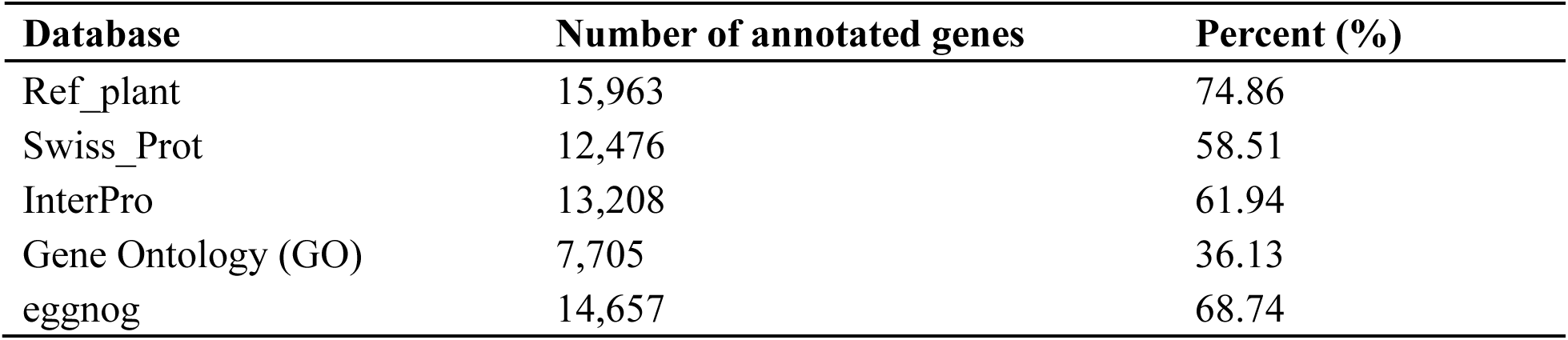
Summary of functional annotations for protein-coding genes in the *Wolffia globosa* genome.

**Table S7.** Summary of repetitive elements in the *Wolffia globosa* genome.

| **Class** | **Subclass** | **Number** | **Length (bp)** | **Percent (%)** |
| --- | --- | --- | --- | --- |
| Retroelements |  | 833,739 | 496,694,207 | 49.59 |
|  | LTR elements | 795,818 | 466,640,181 | 46.59 |
|  | Gypsy | 134,173 | 87,538,006 | 8.74 |
|  | Copia | 193,835 | 104,809,279 | 10.46 |
|  | LINEs | 37,921 | 30,054,026 | 3.00 |
| DNA transposons |  | 604,264 | 196,856,956 | 19.65 |
| Other repeats |  |  |  |  |
|  | Unclassified | 66,587 | 9,029,167 | 0.90 |
| Total |  |  | 702,580,330 | 70.14 |

**Table S9.**
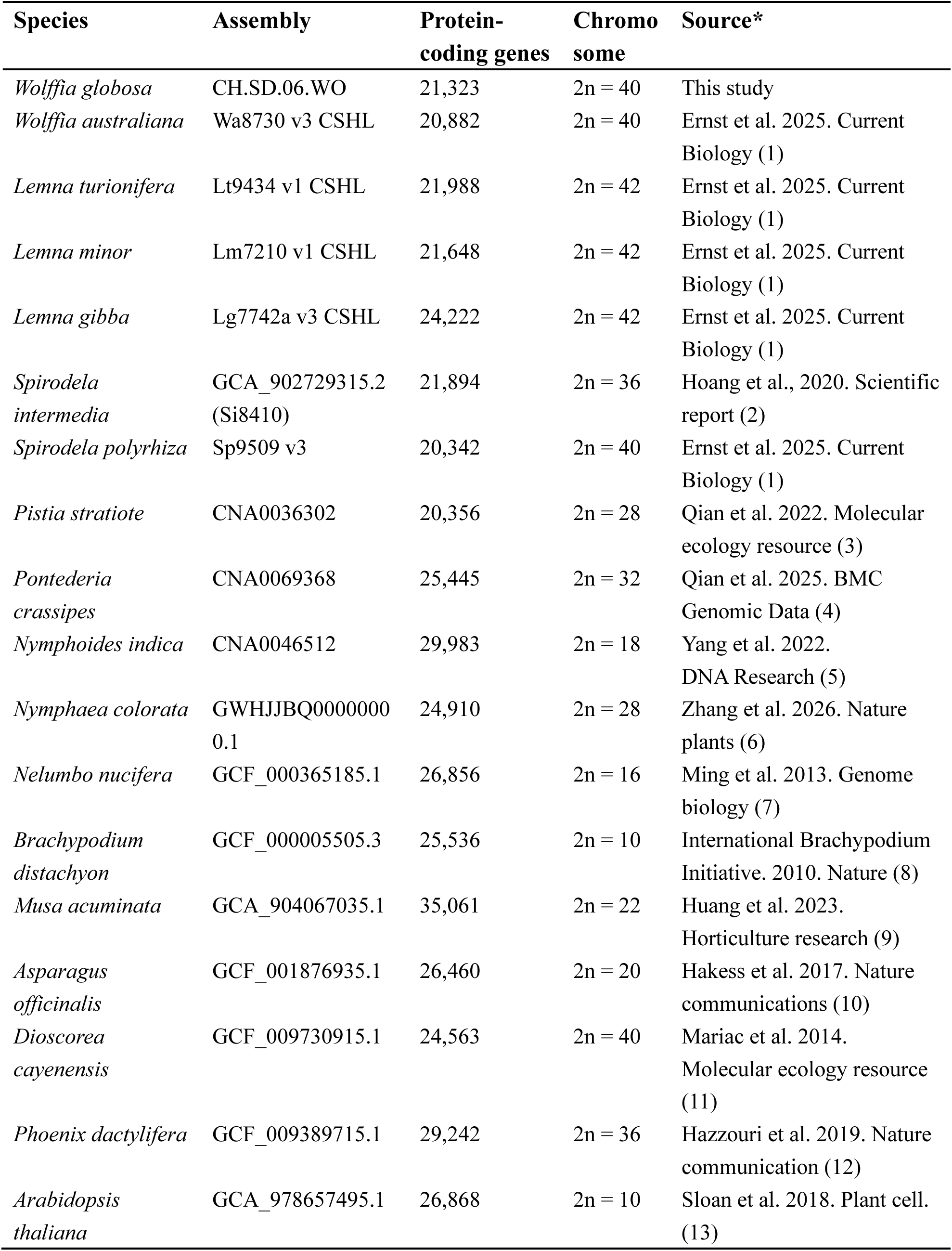
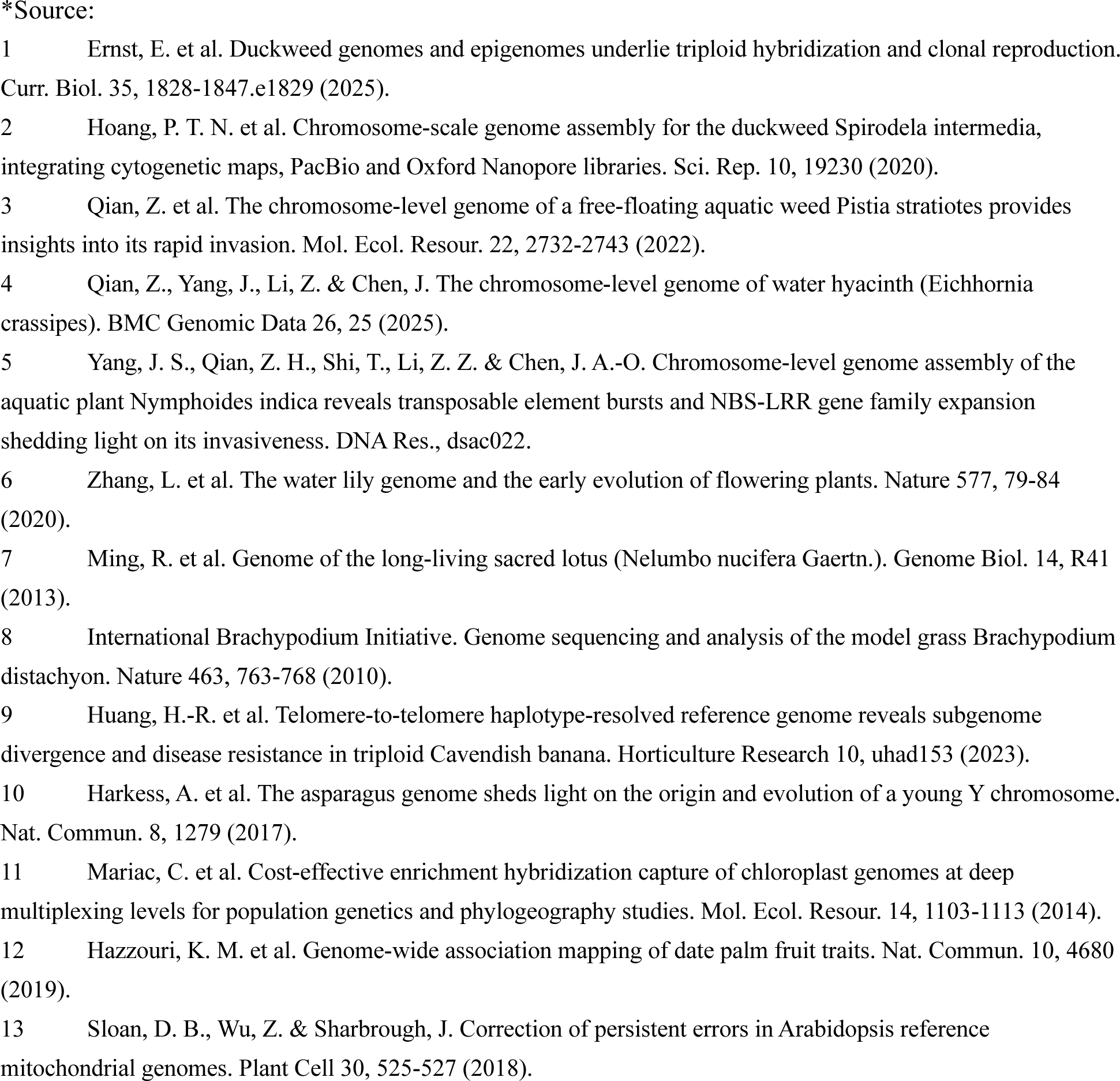
Genome assemblies used in this study.

| Species | Assembly | Protein-coding genes | Chromosome | Source* |
| --- | --- | --- | --- | --- |
| *Wolffia globosa* | CH.SD.06.WO | 21,323 | 2n = 40 | This study |
| *Wolffia australiana* | Wa8730 v3 CSHL | 20,882 | 2n = 40 | Ernst et al. 2025. Current Biology (1) |
| *Lemna turionifera* | Lt9434 v1 CSHL | 21,988 | 2n = 42 | Ernst et al. 2025. Current Biology (1) |
| *Lemna minor* | Lm7210 v1 CSHL | 21,648 | 2n = 42 | Ernst et al. 2025. Current Biology (1) |
| *Lemna gibba* | Lg7742a v3 CSHL | 24,222 | 2n = 42 | Ernst et al. 2025. Current Biology (1) |
| *Spirodela intermedia* | GCA_902729315.2 (Si8410) | 21,894 | 2n = 36 | Hoang et al., 2020. Scientific report (2) |
| *Spirodela polyrhiza* | Sp9509 v3 | 20,342 | 2n = 40 | Ernst et al. 2025. Current Biology (1) |
| *Pistia stratiote* | CNA0036302 | 20,356 | 2n = 28 | Qian et al. 2022. Molecular ecology resource (3) |
| *Pontederia crassipes* | CNA0069368 | 25,445 | 2n = 32 | Qian et al. 2025. BMC Genomic Data (4) |
| *Nymphoides indica* | CNA0046512 | 29,983 | 2n = 18 | Yang et al. 2022. DNA Research (5) |
| *Nymphaea colorata* | GWHJJBQ0000000 0.1 | 24,910 | 2n = 28 | Zhang et al. 2026. Nature plants (6) |
| *Nelumbo nucifera* | GCF_000365185.1 | 26,856 | 2n = 16 | Ming et al. 2013. Genome biology (7) |
| *Brachypodium distachyon* | GCF_000005505.3 | 25,536 | 2n = 10 | International Brachypodium Initiative. 2010. Nature (8) |
| *Musa acuminata* | GCA_904067035.1 | 35,061 | 2n = 22 | Huang et al. 2023. Horticulture research (9) |
| *Asparagus officinalis* | GCF_001876935.1 | 26,460 | 2n = 20 | Hakess et al. 2017. Nature communications (10) |
| *Dioscorea cayenensis* | GCF_009730915.1 | 24,563 | 2n = 40 | Mariac et al. 2014. Molecular ecology resource (11) |
| *Phoenix dactylifera* | GCF_009389715.1 | 29,242 | 2n = 36 | Hazzouri et al. 2019. Nature communication (12) |
| *Arabidopsis thaliana* | GCA_978657495.1 | 26,868 | 2n = 10 | Sloan et al. 2018. Plant cell. (13) |

\*Source:

